# RNA polymerase II is required for spatial chromatin reorganization following exit from mitosis

**DOI:** 10.1101/2020.10.27.356915

**Authors:** Shu Zhang, Nadine Übelmesser, Natasa Josipovic, Giada Forte, Johan A. Slotman, Michael Chiang, Henrike Gothe, Eduardo Gade Gusmao, Christian Becker, Janine Altmüller, Adriaan B. Houtsmuller, Vassilis Roukos, Kerstin S. Wendt, Davide Marenduzzo, Argyris Papantonis

## Abstract

Mammalian chromosomes are three-dimensional entities shaped by converging and opposing forces. Mitotic cell division induces drastic chromosome condensation, but following reentry into the G1 cell cycle phase, condensed chromosomes unwind to reestablish interphase organization. Here, we use a cell line allowing auxin-mediated degradation of RNA polymerase II to test its role in this transition. *In situ* Hi-C showed that RNAPII is required for compartment and loop formation following mitosis. RNAPs often counteract loop extrusion and, in their absence, longer and more prominent loops arise. Evidence from chromatin fractionation, super-resolution imaging and *in silico* modeling attribute these effects to RNAPII-mediated cohesin loading at active promoters upon reentry into G1. Our findings reconcile the role of RNAPII in gene expression with that in chromatin architecture.

## INTRODUCTION

The evolution and expansion of chromosome conformation capture (3C) technologies (Denker and de Laat, 2016; Übelmesser and Papantonis, 2019), has profoundly renewed our understanding of the three-dimensional (3D) organization of eukaryotic chromosomes and how it underlies their function and maintenance (Marchal et al, 2019; Ibrahim and Mundlos, 2020). It is now well accepted that chromosomes are dynamic entities (Hansen et al, 2018), and that their dynamics result from converging and opposing forces acting on chromatin (Rada-Iglesias et al, 2018). These include tethering to nuclear landmarks like the lamina or the nucleolus (Canat et al, 2020), the interplay between transcription factor-bound *cis*-elements (Kim and Shendure, 2019), and the dynamic extrusion of loops via cohesin complexes counteracting the propensity of homotypic compartments to interact with one another (Nuebler et al, 2018; Rada-Iglesias et al, 2018).

Kilobase-resolution Hi-C contact maps revealed thousands of loops genome-wide that are almost invariably anchored at convergent CTCF-bound motifs. These loops and corresponding “loop domains” represent prominent sub-Mbp features of chromosomal organization (Rao et al, 2014). Combining high-resolution Hi-C with auxin-mediated acute and reversible depletion (Yesbolatova et al, 2019) of key chromatin-organizing factors has shed light on their emergence. Depletion of CTCF from mESCs led to loss of insulation at the boundaries of thousands of topologically-associated domains (TADs) (Nora et al, 2017). More strikingly, depletion of cohesin complex subunits led to the elimination of essentially all CTCF-anchored loops (Rao et al, 2017; Schwarzer et al, 2017; Gassler et al, 2017). Depletion of the cohesin release factor WAPL promotes loop enlargement and aberrant loop formation by also engaging non-convergent CTCF anchors (Haarhuis et al, 2017; Gassler et al, 2017). These observations, together with the recently documented ability of cohesin to actively extrude loops *in vitro* (Davidson et al, 2019; Kim et al, 2019) and the finding that CTCF-STAG interactions protect cohesin from chromatin release (Li et al, 2020; Wutz et al, 2020), have crystalized a model for how architectural loops form and dissolve.

In addition to cohesin, another molecular motor known for its ability to translocate DNA is the RNA polymerase (Papantonis and Cook, 2011). However, its contribution to the organization of interphase chromatin folding is still debated. Different lines of evidence point to a connection between RNAPII binding and the differential formation of spatial interactions. To cite recent examples, allele-specific Hi-C showed that the mouse inactive X chromosome lacks active/inactive compartments and TADs, which however form around “escapee” genes and in the active allele (Giorgetti et al, 2016); the transcriptional state of variably-sized domains across eukaryotes, from *C. elegans* and *D. melanogaster* to *A. thaliana* and mammals, is a robust predictor of interactions mapped via Hi-C and explains chromatin partitioning to a great extent (Ulianov et al, 2016; Rowley et al, 2017); and TAD emergence coincides with the activation of transcription in the zygote (Hug et al, 2017). Abrogation of transcription using inhibitors weakens, but does not alleviate, TAD boundary insulation (Hug et al, 2017; Barutcu et al, 2019), while RNase treatment of native or fixed nuclei does not affect TADs, but eliminates specific contacts (Brant et al, 2016; Barutcu et al, 2019). Single-nucleosome imaging upon rapid RNAPII depletion showed that polymerases act to constrain and direct chromatin movement in 3D space, compatible with the idea of a transcription-based chromatin organization in “factories” or “condensates” (Nagashima et al, 2019).

On the other hand, RNAPII and Mediator-complex components were found to be dispensable for bringing *cis*-elements into spatial proximity (El Khattabi et al, 2019), and pharmacological inhibition of transcription in parallel with RAD21 reintroduction in cohesin-depleted cells did not affect CTCF loop reestablishment (Rao et al, 2017). At the same time, CTCF or cohesin depletion from mammalian cells had rather limited impact on gene expression (Nora et al, 2017; Rao et al, 2017), and upon CTCF loop elimination, a comparable number of loops formed on the basis of chromatin identity (Rao et al, 2017) or did not dissolve at all (Thiecke et al, 2020). Most recently, Micro-C, a sub-kbp Hi-C variant, unveiled thousands of fine-scale loops connecting transcriptionally-active loci in mouse and human cells, often without association to CTCF/cohesin (Krietenstein et al, 2020; Hsieh et al, 2020).

On top of its potentially direct effects, RNAPs and the act of transcription remodel 3D genome folding via interplay with cohesin-CTCF complexes. For example, transcription can relocate cohesin by many kilobases (Busslinger et al, 2017). Such transcription-mediated displacement can even disrupt prominent CTCF loops and rewire spatial interactions (Heinz et al, 2018). In addition, in an antagonistic relationship with condensin complexes, RNAPs are essential for domain formation and opposed by condensin (Brandão et al, 2019; Rowley et al, 2019). This and other data highlight the need to dissect and reconcile RNAPII contributions of in chromatin organization. To this end, and as pharmacological inhibition of RNAPs is inefficient, we exploited a human cell line that allows for the rapid and reversible RNAPII depletion from cells (Nagashima et al, 2019). We combined *in situ* Hi-C and super-resolution imaging in the presence or absence of RNAPII with *in silico* models to disentangle its role in gene expression from that in genome architecture.

## RESULTS

### Acute RNAPII depletion impacts loop-level interphase folding

RNAPII is essential for cell viability, so its depletion may only be transient. We exploited a human DLD-1 colorectal cancer line, in which the larger RNAPII subunit, RPB1, is tagged with a mini-AID domain and can, thus, be acutely and reversibly degraded upon addition of auxin (and doxycycline to activate the plant ubiquitin ligase TIR1 recognizing the mAID domain; Nagashima et al, 2019). In our hands, 2 hours of dox/auxin treatment reduce RNAPII protein levels by >60%, while 14 h of treatment result in >90% RNAPII degradation without affecting RNAPI or III levels (**Figures 1A** and **S1A**). Degradation is somewhat less impactful for RNAPs phosphorylated at Ser5 residues in their C-terminal domain repeats (CTD; Heidemann and Eick, 2012). Washing out auxin in the presence of its competitive inhibitor, auxinole, largely restores RNAPII-Ser5 levels (**Figure S1A**), suggesting that soluble RNAPII is the most susceptible to degradation and that essentially all polymerases restored upon auxin removal are directly embedded into chromatin.

**Figure 1.**
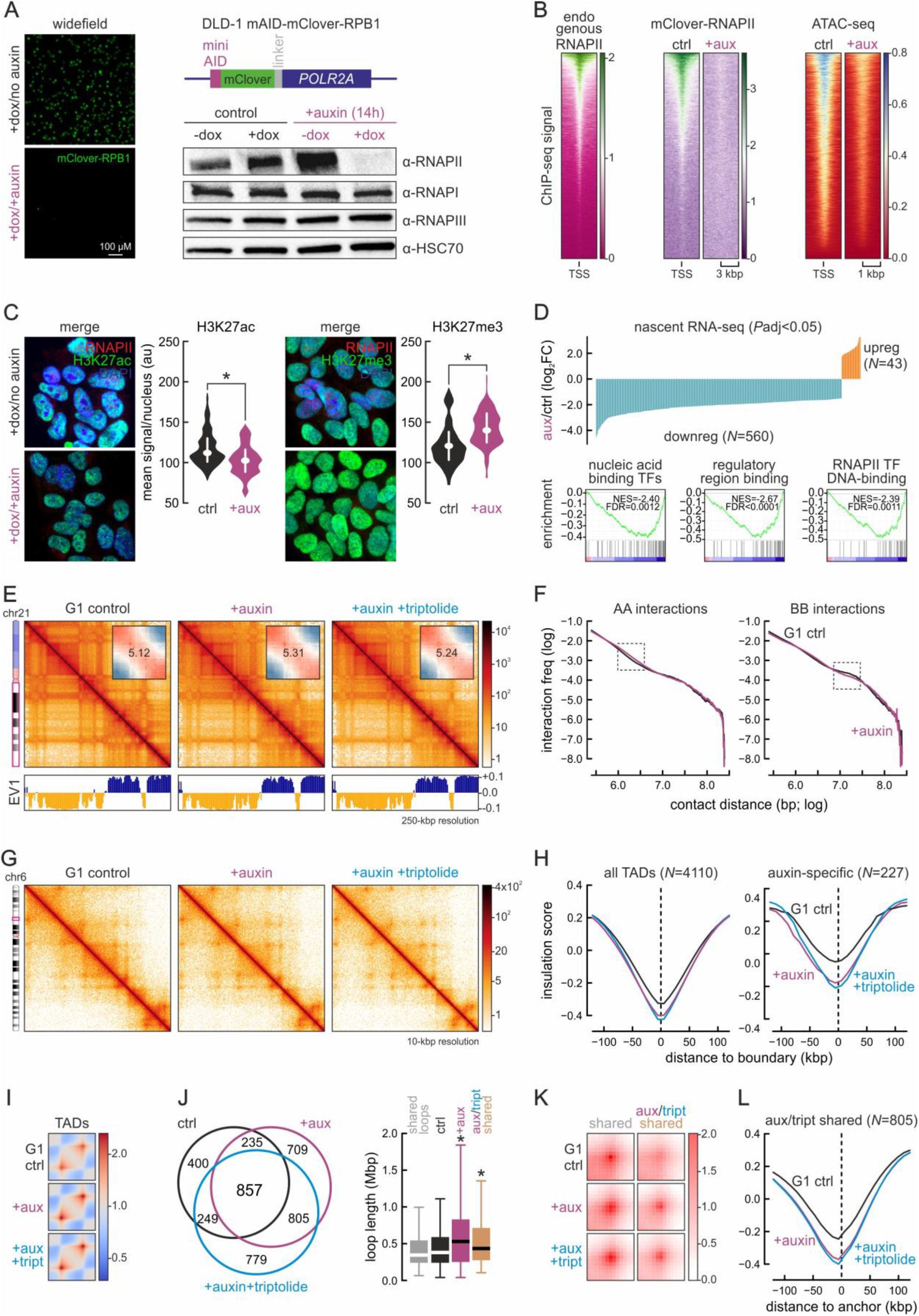
Effects of RNAPII acute degradation on interphase chromatin folding. (**A**) *Left*: DLD1-mAID-RPB1 cells lose mClover signal fused to the large RNAPII subunit (schematic) upon doxycycline/auxin treatment for 14 h. *Right*: Selective RNAPII degradation confirmed by Western blotting; HSC70 levels provide a loading control. (**B**) Heatmaps showing mClover-RNAPII ChIP-seq signal overlapping positions bound by RNAPII in parental DLD-1 (GSM2769059) and its loss upon auxin treatment (left/middle) concomitant with a decrease in ATAC-seq signal (right). (**C**) Representative H3K27ac and H3K27me3 immunofluorescence from untreated (top) or 14-h auxin-treated cells (bottom) and signal quantification (bean plots). *: significantly different; *P*<0.01, Wilcoxon-Mann-Whitney test. Bar: 5 μM. (**D**) *Top*: Graph showing nascent RNA changes (log_2_ fold-change compared to control cells, *P*_adj_<0.05) in 603 genes upon auxin treatment. *Bottom*: Gene set enrichment analysis. (**E**) Exemplary Hi-C maps of chr21 from G1-sorted control (left), 14-h auxin-treated (middle), and auxin plus triptolide-treated cells (right) at 250-kbp resolution aligned to first eigenvector values (below). *Insets*: saddle plots showing A/B-compartment insulation. (**F**) Decay plots showing Hi-C interaction frequency amongst A- (left) or B-compartments (right) as a function of genomic distance (log) in control (black line) or auxin-treated cells (purple line). Dashed rectangles indicate distances at which the two lines deviate the most. (**G**) Exemplary Hi-C maps of chr6 from control (left), auxin-treated (middle), and auxin plus triptolide-treated cells (right) at 10-kbp resolution. (**H**) Line plots showing mean insulation scores from control (black line), auxin-treated (purple line), and auxin plus triptolide-treated cells (blue line) in the 240 kbp around all (left) or degron-specific TAD boundaries (right). The number of TADs queried (*N*) is indicated. (**I**) Heatmaps showing mean TAD-level interactions in control (top), auxin-treated (middle), and auxin-/triptolide-treated cells (bottom). (**J**) *Left*: Venn diagram showing shared and unique loops to control (black), auxin-treated (purple), and auxin plus triptolide-treated Hi-C (blue line). *Right*: Loop lengths displayed as boxplots (right). *: significantly different; *P*<0.01, Wilcoxon-Mann-Whitney test. (**K**) APA plots showing mean Hi-C signal for the different loop groups in panel I. (**L**) As in panel H, but for the loop anchor shared only by auxin-treated and auxin plus triptolide-treated cells.

To further characterize this line, we performed RNAPII ChIP-seq using an antibody targeting the mClover tag in RBP1. When compared to public ChIP-seq data of DLD-1 RNAPII (Chen et al, 2017), mClover-tagged polymerases occupied the same positions and were fully removed from chromatin upon auxin treatment (**Figure 1B**). Polymerase degradation was accompanied by a strong decrease in chromatin accessibility at TSSs genome-wide (**Figure 1B**), similar to what was recently observed using mESCs (Jiang et al, 2020). We also queried the H3K27ac (denoting active chromatin) and H3K27me3 histone marks (marking facultative heterochromatin) upon auxin treatment. We observed significant H3K27ac reduction concomitant with increased H3K27me3 levels (**Figure 1C**). Finally, we monitored changes in nascent RNA production using “factory” RNA-seq (Caudron-Herger et al, 2015). Control and auxin-treated samples separated well in PCA plots (**Figure S1B**) and, focusing on strongly deregulated genes (i.e., log_2_FC of at least ±2, *P*_adj_<0.05), we found >600 changing significantly. Of these, >90% were downregulated and mainly involved in transcription, chromatin binding, and cell cycle regulation (**Figures 1D** and **S1B**).

Given these robust effects, we asked whether the spatial organization of interphase chromatin is also altered upon RNAPII removal. We applied *in situ* Hi-C to G1-sorted DLD1-RPB1-mAID cells treated or not with auxin for 14 h or to cells in which auxin treatment was complemented by triptolide, a TFIIH inhibitor abrogating transcriptional initiation and further enhancing RNAPII degradation (Wang et al, 2011). The use of G1 cells removes heterogeneity coming from S-/G2-phase cells to generate Hi-C maps of higher detail (Sati et al, 2020). Following data analysis, we saw marginal differences at the level of A- /B-compartments (**Figure 1E,F**). At higher resolution, TADs also only showed mild disruptions (**Figure 1G**), with few TADs (<20%) of the 4110 identified in control Hi-C data changing in cells lacking RNAPII. Natably, >200 TADs appeared *de novo* in auxin-/triptolide-treated cells and displayed reinforced insulation at their boundaries (**Figure 1H**). Moreover, average Hi-C profiles in/around TADs, revealed increased definition of their borders at the expense of intra-TAD interactions (**Figure 1I**).

Finally, >1500 loops formed exclusively in auxin-treated cells. These were significantly larger than those in control cells or those shared between conditions (**Figure 1J,K**). Of these, 805 loops shared by the auxin- and auxin-/triptolide-treated cells were also larger and displayed increased insulation at their anchors (**Figure 1J-L**). Note that same analyses on Hi-C data from mAID-RBP1 cells treated with auxin for 2 h (where 60-70% RPB1 is degraded; **Figure S1A**), did not reveal changes at any level of 3D genome organization (**Figure S1C-K**). Together, our findings argue that subtle yet discernible effects on TAD- and loop-level organization occur upon strong RNAPII depletion, and come to add to recent data, where RNAPII depletion in asynchronous mouse ESCs led to mild changes in folding (Jiang et al, 2020).

### Reestablishment of spatial chromatin organization after mitosis requires RNAPII

Given that RNAPII degradation did not severely affect interphase chromatin organization, we tested whether it is implicated in reestablishing chromatin folding upon exit from mitosis. This was based on two observations. First, on the detailed description of chromatin refolding dynamics in the mitosis-to-G1 transition, where contacts among *cis*-elements form early and rapidly, and are often not related to CTCF/cohesin (Abramo et al, 2019; Zhang et al, 2019). Second, on the fact that, early in this transition, >50% of all active genes and enhancers exhibit a strong transcription spike (Hsiung et al, 2016).

To study chromatin refolding following mitotic exit, we synchronized mAID-RPB1 cells at the G2/M checkpoint using the CDK1 inhibitor RO3306 (blocking ^~^90% cells in G2; **Figure 2A,B**), before releasing them via mitosis into G1 by washing out the inhibitor for 6.5 h. This allowed >70% cells to reenter G1 and be collected by FACS (**Figure 2A,B**). RNAPII degradation was initiated by adding auxin to cells while arrested in G2, was maintained throughout mitosis and G1 reentry without compromising progression past early G1. Strong degradation was confirmed by chromatin fractionation blots (**Figure 2C**) and EUTP-labeling of nascent RNA (**Figure S2A**). Like in asynchronous cells, H3K27ac levels were decreased and H3K27me3 levels increased in G1-reentry cells depleted of RNAPII (**Figures 2C** and **S2B-D**). At the same time, the presence of abundant SWI/SNF chromatin remodeler subunits in chromatin was affected, but CTCF incorporation remained largely unchanged (**Figure 2C**).

**Figure 2.**
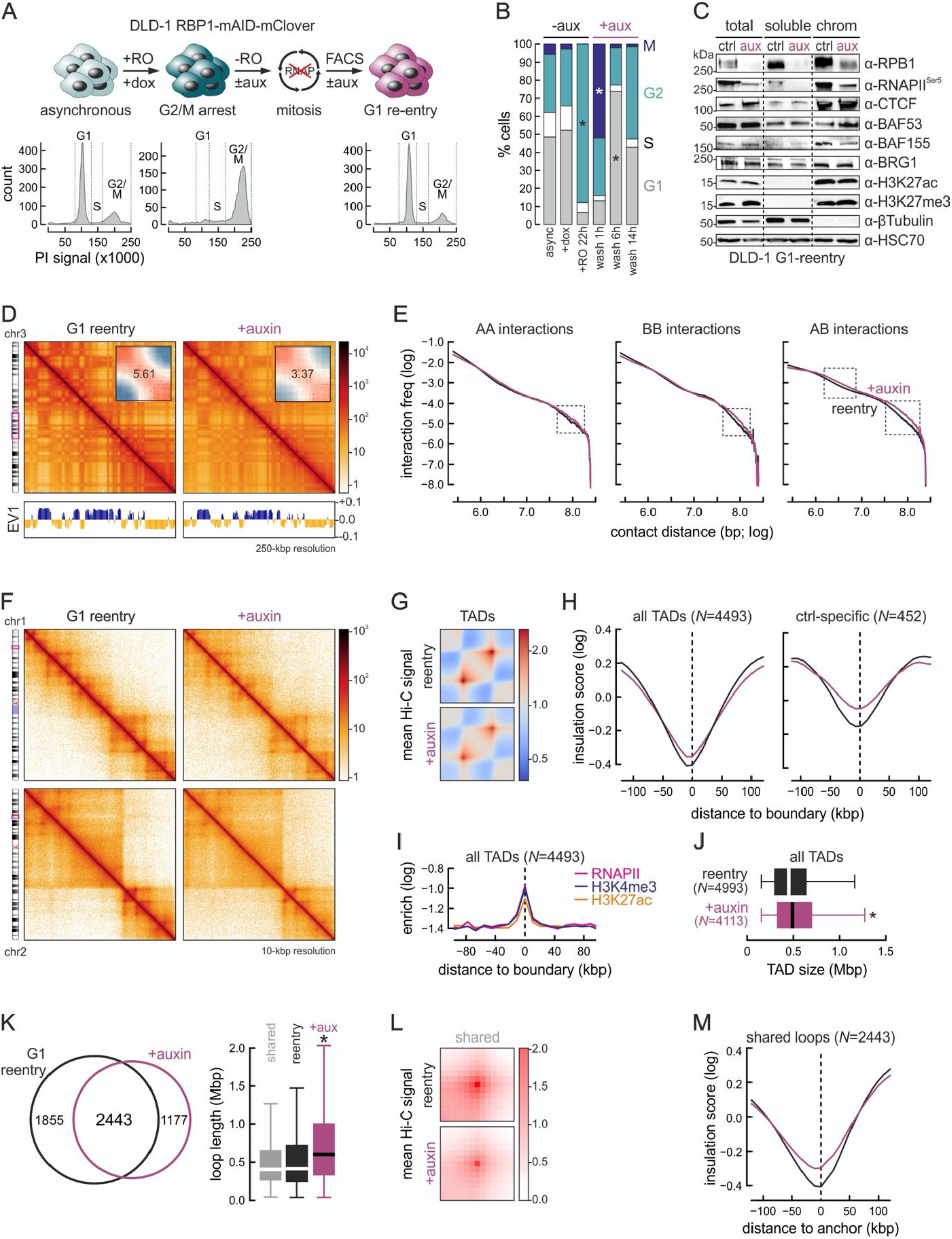
RNAPII contributes to genome refolding following exit from mitosis. (**A**) *Top*: Overview of the experimental scheme for DLD1-mAID-RBP1 cell synchronization. *Bottom*: Propidium iodine FACS profiles at each step. (**B**) Bar plots quantifying the percent of cells in each cell cycle phase from the procedure in panel A. *: significantly different; *P*<0.01, Fischer’s exact test. (**C**) Fractionation western blots showing changes in chromatin-bound RNAPII, chromatin remodelers and histone mark levels; HSC70 provides a control. (**D**) Exemplary Hi-C maps of chr3 from control (left) and auxin-treated reentry cells (right) at 250-kbp resolution aligned first eigenvector values (below). *Insets*: saddle plots showing loss of A/B- compartment insulation. (**E**) Decay plots showing Hi-C interaction frequency between A- (left), B- (middle) or A/B-compartments (right) as a function of genomic distance (log) in control (black line) and auxin-treated reentry cells (purple line). Dashed rectangles indicate distances where the two lines deviate the most. (**F**) Exemplary Hi-C maps of chr1 and 2 from control (left) and auxin-treated reentry cells (right) at 10-kbp resolution. (**G**) Heatmaps showing mean TAD-level interactions in control (top) and auxin-treated reentry cells (bottom). (**H**) Line plots showing mean insulation scores from control (black line) and auxin-treated reentry cells (purple line) in the 240 kbp around all (left) or control-specific TAD boundaries (right). The number of TAD boundaries queried (*N*) is indicated. (**I**) Line plot showing RNAPII (magenta; GSM2769059), H3K4me3 (blue; GSM2283764) and H3K27ac ChIP-seq signal enrichment (orange; GSM2037784) in the 100 kbp around TAD boundaries from control cells. (**J**) Boxplots showing TAD sizes in control (black) and auxin-treated reentry cells (purple). *: significantly different; *P*<0.01, Wilcoxon-Mann-Whitney test. (**K**) *Left*: Venn diagram showing shared and unique loops between control (black) and auxin-treated reentry cells. *Right*: Loop lengths are displayed as boxplots. *: significantly different; *P*<0.01, Wilcoxon-Mann-Whitney test. (**L**) APA plots showing decreasing Hi-C signal for shared loops from panel J. (**M**) As in panel H, but for loop anchors shared between control and auxin-treated reentry cells.

We next performed Hi-C on G1-reentry cells treated or not with auxin. Libraries were sequenced to >1 billion read pairs, and contained >740 million and >1 billion Hi-C contacts for control and auxin-treated samples, respectively (**Table S1**). Our first observation was that RNAPII-depleted cells showed increased interchromosomal contacts developing at the expense of intrachromosomal ones (**Figure S2E,F**). This was orthogonally confirmed using high throughput 3D-DNA FISH (**Figure S2G**). At the same time, compartment boundaries in *cis* were markedly blurred (**Figures 2D** and **S2H**), with interactions between A- and B-compartment segments becoming stronger at distances >1 Mbp (**Figure 2E**).

Loss of interactions at the TAD scale (<1 Mbp) in auxin-treated cells (**Figure S2I**) led us to analyze 10-kbp resolution Hi-C maps. There, we observed strong and widespread erosion of domain structure, local insulation and loop formation (**Figures 2F,G** and **S2J**). Insulation was weakened across all ^~^4500 TADs identified in control reentry cells, and markedly more in the ^~^10% of TAD boundaries that could not be reestablished in the absence of RNAPII (**Figure 2H**). These effects are in line with RNAPII and active histone mark enrichment at TAD boundaries (**Figure 2I**). Overall, RNAP-depleted reentry cells had fewer and larger TADs than control cells (**Figure 2J**), indicative of boundary collapse and TAD merging.

At the loop level, ^~^1900 were lost from RNAPII-depleted cells concomitantly with the emergence >1150 new, significantly longer loops (**Figure 2K**). Curiously, loops occurred *de novo* at sites of existing insulation, whereas loops not reestablished in RNAP-depleted cells displayed reduced insulation (**Figure S2K,L**). The 2443 loops detected in both control and auxin-treated reentry cells were weaker in the absence of RNAPII while also displaying reduced insulation at their anchors (**Figure 2K-M**). Last, looking at the formation of “stripes”, considered a readout of loop extrusion (Vian et al, 2018), they were both accentuated and dissolved in RNAPII-depleted cells (**Figure S2M**). In summary, our data suggest that RNAPII is critically implicated in reestablishing both higher-order and fine-scale chromatin folding upon exit from mitosis, and its depletion compromises loop formation. Also, the folding changes that follow polymerase depletion do not simply reflect structure of an early G1 time-point (when compared to data from Zhang et al, 2019), but rather compromised refolding.

### Depleting topoisomerase II does not affect chromatin folding upon G1-reentry

Transcriptional elongation enforces supercoiling onto DNA and topoisomerase I (TOP1) was recently shown to be stimulated by RNAPII to resolve supercoiling as elongation progresses. However, TOP1 binding at TSSs alongside initiating polymerases was not matched by potent TOP1 activity (Baranello et al, 2016). On the other hand, TOP2 has been linked to chromatin organization along the cell cycle and to transcription (Nitiss, 2009; Bunch et al, 2015) with TOP2A affecting RNAPII kinetics (Thakurela et al, 2013) and marking RNAPII pausing sites (Singh et al, 2020). Moreover, TOP2B flanks TAD boundaries in human cells alongside CTCF/cohesin complexes (Uusküla-Reimand et al, 2016), and it confines RNAPII and preserves domain boundaries in yeast (Achar et al, 2020).

Given that no elongating RNAPII remains in auxin-treated cells, and that TOP2A-mAID cells prolong, but do conclude mitosis (Nielsen et al, 2020), we asked whether TOP2 depletion from G1-reentry cells explains the effects we observe in RNAPII-depleted cells. To this end, we exploited another colorectal cancer line, HCT116, carrying a full knockout of the *TOP2B* gene and expressing mAID-tagged TOP2A. We verified >80% depletion of TOP2A, and used the same synchronization and FACS sorting scheme as before to obtain G1-reentry cells (**Figure S3A,B**). Our first observation, using “factory” RNA-seq, was that only 155 genes were affected by TOP2 elimination (log_2_FC at least ±0.6, *P*_adj_<0.01), and these were linked to cell cycle control and DNA-binding complexes (**Figure S3C**).

Hi-C performed on G1-reentry cells treated or not with auxin revealed marginal changes across all scales of chromatin organization. Compartments were not affected, interactions remained unchanged irrespective of distance, and no increase in *trans* contacts was seen (**Figure S3D-F**). Negligible changes to TAD boundary insulation were observed, and the mean size of TOP2A/B-depleted TADs did not differ from that of control cells (**Figure S3G-J**). Finally, at the level of loops, ^~^950 and ^~^850 loops were lost or gained upon TOP2-depletion, respectively, and condition-specific loops were again larger (**Figure S3K**). However, the increase/reduction in Hi-C signal at these loops was significantly less than that recorded upon RNAPII depletion, and not followed by changes in insulation at the respective loop anchors (**Figure S3L,M**). In summary, these data suggest that the effects inflicted on chromatin refolding by RNAPII degradation are not due to compromised TOP2A/B activity, but must rather be polymerase-centric.

### RNAPII removal compromises cohesin reloading onto chromatin and loop extrusion

Hi-C data from our G1-reentry cells lacking RNAPII clearly demonstrate A/B-compartment mixing and differential loss/gain of loops and loop domains. These effects are reminiscent of those seen in RAD21-mAID (Rao et al, 2017) or *Nipbl*-knockout cells (Schwarzer et al, 2017). To reconcile these observations, we examined how the levels of cohesin subunits change in reentry cells following auxin treatment. First, fractionation western blots showed little fluctuation in SMC1A or RAD21 levels in chromatin, which was confirmed by RAD21 and NIPBL quantification in single cells via immunofluorescence (**Figure 3A,B**). At the same time, the two cohesin loaders, NIPBL and MAU2, were markedly reduced in the chromatin fraction, as were the levels of the protein responsible for cohesin unloading, WAPL (concomitant with an increased in soluble pool titers; **Figure 3A**).

**Figure 3.**
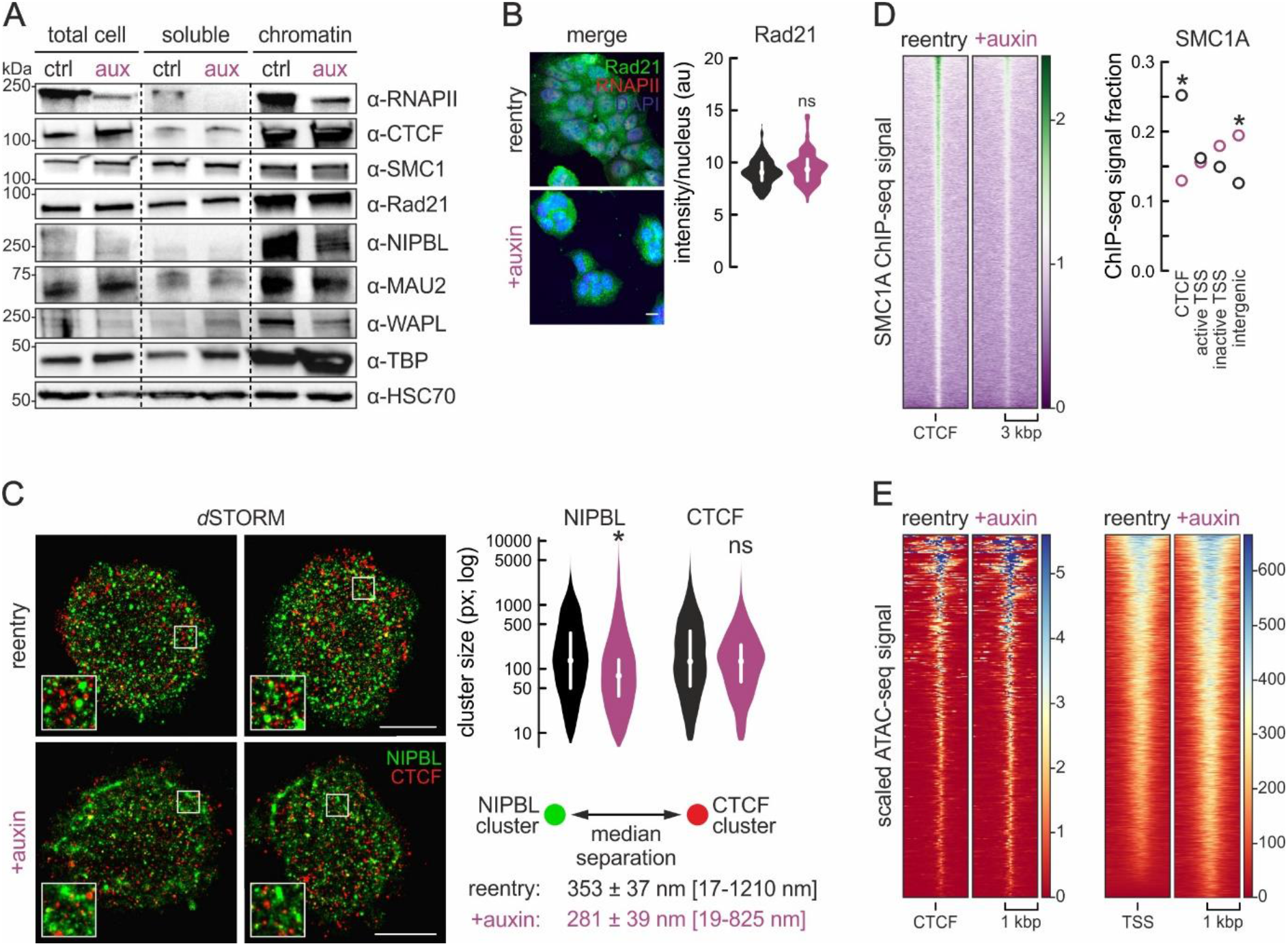
RNAPII degradation compromises cohesin chromatin reloading upon exit from mitosis. (**A**) Fractionation blots showing changes in chromatin-bound RNAPII, cohesin loaders NIPBL and MAU2, the WAPL unloader, and TBP; HSC70 provides a loading control. (**B**) RAD21 and RNAPII immunofluorescence in untreated (top) or auxin-treated reentry cells (bottom) and signal quantification (bean plots). Bar: 5 μM. (**C**) *Left*: Rendering of 3D-STORM localizations for NIPBL and CTCF from control (top row) and auxin-treated reentry cells (bottom row). Bar: 5 μM. *Right*: Bean plots showing changes in NIPBL and CTCF cluster sizes. *Bottom right*: Changes in separation between the nearest NIPBL/CTCF clusters (smallest and largest distances shown in square brackets). *: significantly different; *P*<0.01, Wilcoxon-Mann-Whitney test. (**D**) *Left*: Heatmaps showing CTCF-proximal SMC1A ChIP-seq signal in auxin-treated versus control reentry cells. *Right*: Plot showing the fraction of normalized SMC1A signal from control (black circles) or auxin-treated ChIP-seq (purple circles) assigned to CTCF-bound, active/inactive TSS or intergenic regions. *: significantly different; *P*<0.01, Fischer’s exact test. (**E**) As in panel D, but using ATAC-seq around CTCF-proximal SMC1A-bound positions (left) or all active TSS (right).

To corroborate these findings, we performed super-resolution localizations of NIPBL and CTCF. Dual-color *d*STORM in control and auxin-treated reentry cells led to the following observations. First, NIPBL localizes in clusters of smaller average size upon RNAPII depletion. Despite this, we also observed more localizations into extended deformed clusters (**Figure 3C**), exemplified by the shift in eccentricity of NIPBL clusters from 5.4 in control to 6.9 in auxin-treated cells (eccentricity of 0 refers to a perfect circle, while eccentricity of 1 to a line). Second, CTCF clusters do not change as regards their mean size, but 50% of all CTCF clusters in control reentry cells lie <129 px^2^, while in auxin-treated cells 50% lie <82 px^2^ (**Figure 3C**). Such an increase in the population of smaller CTCF clusters was also observed using *d*STORM upon cohesin removal from RAD21-mAID cells (Casa et al, 2020) and points to loop collapse. Third, NIPBL distribution relative to CTCF clusters also changed significantly in the absence of RNAPII. The median separation between NIPBL and its nearest CTCF cluster was reduced from 353 to 281 nm, with the largest recorded distance dropping from >1200 to 825 nm (**Figure 3C**).

These results, and NIPBL/MAU2/WAPL protein levels, suggest aberrant cohesin loading to (and most likely unloading from) chromatin in the absence of RNAPII, and predict that less cohesin will end up at CTCF-bound sites, despite its roughly unchanged levels in chromatin. We generated SMC1A ChIP-seq in control and auxin-treated reentry cells and indeed found cohesin signal reduced at CTCF sites genome-wide (**Figure 3D**). We quantified the fraction of normalized ChIP-seq signal falling into CTCF-bound regions, active or inactive TSSs, and non-RNAPII-associated intergenic space. A larger fraction of signal mapped to intergenic regions and inactive TSSs in auxin-treated compared to control cells, while the significant drop is signal fraction at CTCF sites was confirmed (**Figure 3D**). These changes occurred in spite of the unchanged accessibility of these CTCF-proximal positions as judged by ATAC-seq (**Figure 3E**). Finally, cohesin loading might occur at TSSs simply because they are rendered accessible due by RNAPII. Hence, reduced accessibility would readily explain compromised loading. To our surprise, ATAC-seq signal in RNAP-depleted TSSs rather increased, as were TBP levels in chromatin (**Figure 3A,E**). This argues in favor of RNAPs recruiting cohesin to these sites, while setting up TSS nucleosome architecture in G1-reentry cells likely relies on factors preceding the polymerase.

### Computational modeling dissects the role of RNA polymerases in loop extrusion

To dissect the connection between RNAPII and cohesin reloading onto chromatin, we turned to *in silico* modeling of chromatin folding. This allowed us to test scenarios that would be challenging to address experimentally. First, we performed 3D chromatin folding simulations using the HiP-HoP model (Buckle et al, 2018) that accounts for the heteromorphic nature of chromatin, and incorporates transcription factor binding (Brackley et al, 2016) and loop extrusion (Fudenberg et al, 2016). We modeled a 10-Mbp region from HUVEC chr14 for which data on gene expression, histone marks, and CTCF positioning are available (ch14:50-60 Mbp, hg19; www.encodeproject.org).

Wild-type chromatin folding was simulated by assuming that most cohesin loading (90%) occurs at RNAPII-occupied TSSs (with 10% loading randomly). Experimentally-defined cohesin residence times on DNA (^~^20 min; Gerlich et al, 2006; Hansen et al, 2017) was incorporated into the model. Following multiple iterations, our model produced a mean contact map resembling Hi-C data (Zirkel et al, 2018; Buckle et al, 2018). To simulate the folding of chromatin in the context of strong RNAPII degradation, we eliminated loading at promoters, and only allowed for random loading (consistent with low efficacy NIPBL-independent cohesin loading *in vitro*; Davidson et al, 2016). As a result of this, four major effects were observed. First, a general weakening of interactions and domain insulation across the 10 Mbp modeled (**Figure 4A**), similar to what we saw using Hi-C (**Figure 2F**). Second, individual models of the fiber showed obvious unfolding (**Figure 4B**), probably consistent with the increase in *trans* interactions in our data (**Figure S2D,E**). Third, reduced cohesin occupancy at CTCF sites (**Figure 4A**), consistent with our SMC1A ChIP-seq (**Figure 3D**). Fourth, markedly weakened loop formation, but with larger loop sizes (**Figure 4D**) closely matching our experimental results (**Figure 2K,L**). Thus, these modeling data suggest that inability to load cohesin at RNAPII-bound sites suffices for explaining chromatin folding differences observed experimentally.

**Figure 4.**
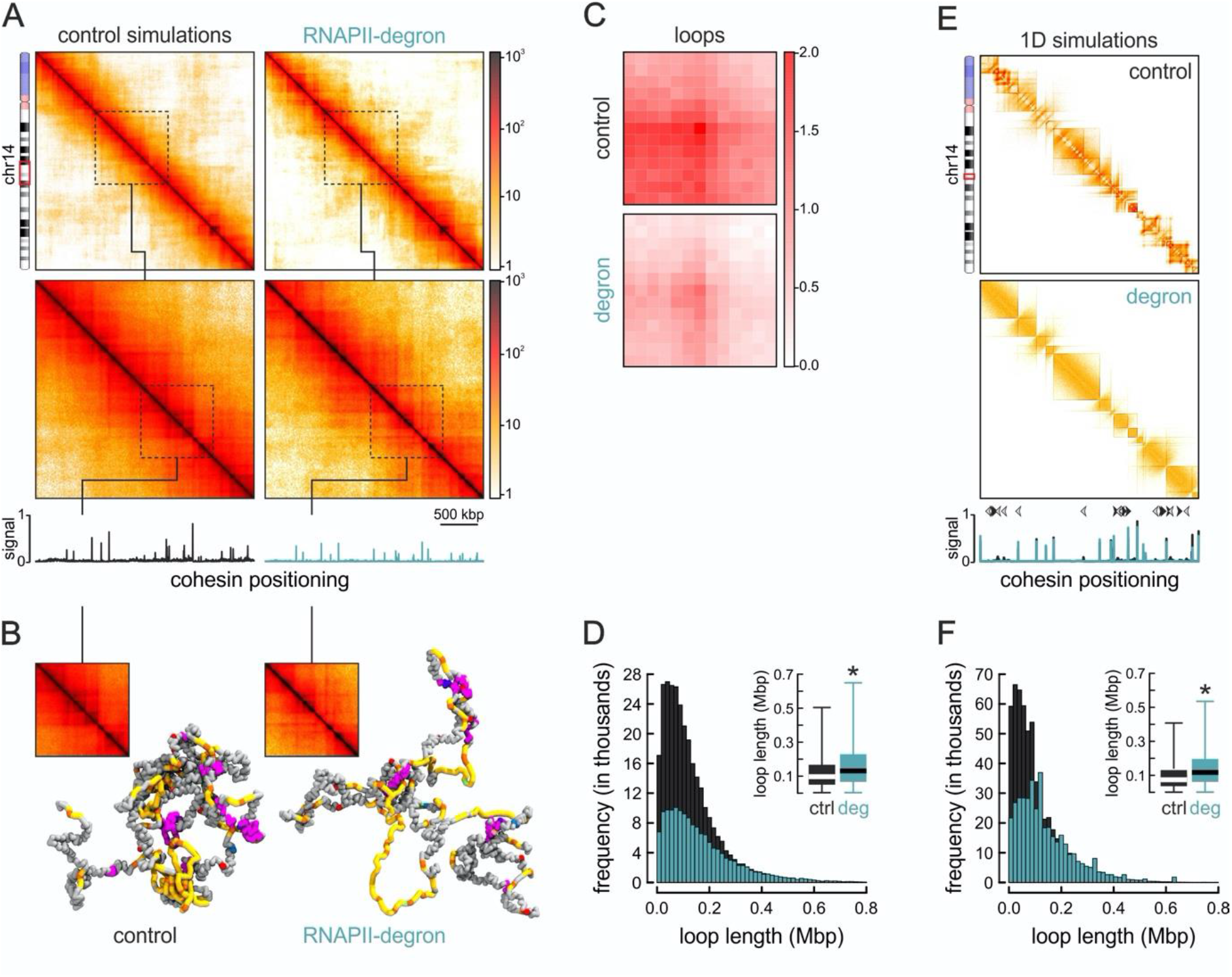
Modeling effects of RNAPII degradation on cohesin loading. (**A**) *Top*: Heatmaps rendered from simulations of wild-type (left) or RNAPII-depleted models (right) of HUVEC chr14:50-60 Mbp. *Bottom*: 10 kbp-resolution heatmaps in the chr14:53-56 Mbp region. Cohesin positioning tracks are aligned below each heatmap. (**B**) Exemplary 3D chromatin folding models of the chr14: 54.5-55.5 Mbp subregion. (**C**) APA plots showing weakened loops in RNAPII-depleted models. (**D**) Histogram showing looping frequency in the absence of RNAPII (turquoise) compared to wild-type models (black). *Inset*: Boxplots showing larger loop lengths in RNAPII-depleted models (blue). *: significantly different; *P*<0.01, Wilcoxon-Mann-Whitney test. (**E**) Heatmaps rendered from 1D simulations representing wild-type (left) or RNAPII-depleted models (right) of the chr14 segment from panel A at 3-kbp resolution. Cohesin positioning (overlaid tracks) and TSS orientation (arrows) are aligned below. (**F**) As in panel D, but using data from the 1D simulations in panel E.

To interrogate the interplay between RNAPII and loop-extruding cohesins directly, we performed 1D simulations of minimal composition. We modeled a 3-Mbp region of HUVEC chr14 (ch14:53-56 Mbp, hg19) as a coarse-grained fiber carrying CTCF at the appropriate positions, as well as RNAPs transcribing genes in the correct orientation. As before, cohesin was predominantly loaded at RNAPII-bound TSSs in the control scenario, but only randomly in the “degron” model. First, we observed formation of loops and intricate domain compartmentalization under control settings, despite only two activities operating on the fiber (**Figure 4E**). Notably, transcription does affect cohesin deposition and loop formation our model, as exemplified by simulations in which all genes in this 3-Mbp segment were modeled as being transcribed in tandem (**Figure S4**). RNAP depletion eliminated compartmentalization and the frequency of looping was again drastically reduced (**Figure 4E,F**). Cohesin occupancy was decreased at most loop anchors, with loops again becoming larger (**Figure 4E,F**). This parsimonious model allows us to deduce that the effects observed in experiments can, for their most part, be explained by a simple relationship between cohesin loading at RNAP-occupied sites and by the interplay between active transcription and loop extrusion.

## DISCUSSION

Following cell division, mitotic chromosomes unwind in order to reestablish interphase folding. Cohesin is not present on mitotic chromatin and needs to be reloaded. Its reloading coincides with the extrusion of CTCF-anchored loops and TAD reemergence (Abramo et al, 2019; Zhang et al, 2019). However, A/B-compartments, driven by homotypic chromatin interactions, reestablish more rapidly, as do contacts amongst *cis*-regulatory elements. Interestingly, the latter display rates that exceed those of extruded loops (Zhang et al, 2019). In parallel, the general transcription factor TBP bookmarks mitotic chromatin to facilitate gene reactivation (Teves et al, 2018), and as transcription reinitiates in late telophase, a strong activity spike occurs in most genes and enhancers (Hsiung et al, 2016). These observations suggest that RNAP activity may play a central role in reestablishing interphase chromatin organization.

Here, using an RPB1 “degron”, we show that RNAPII presence on chromatin is necessary for both the establishment of compartments and the extrusion of loops. The former is intuitively justified by the homotypic interactions that build the “active” A-compartment, and the recent finding that chromatin acetylation itself has a propensity for compartment formation (Rosencrance et al, 2020; Gryder et al, 2019). In our RNAPII-depleted G1-reentry cells, H3K27ac levels are reduced, H3K27me3 levels increase, and this imbalance most probably underlies compartment-level changes.

The latter effect is more perplexing, but agrees with NIPBL binding active gene promoters before cohesin does following mitotic exit (Zuin et al, 2014; Busslinger et al, 2017) – and, thus, raises three key questions. First, how are compartments and loop domains affected, but most TADs unchanged? Indeed, we only saw a weakening of boundaries in ^~^10% of TADs genome-wide, which led to domain merging. Nonetheless, there was an overall loss of boundary strength, and intra-TAD interactions weakened. Such finer scale changes are in line with HiChIP data in RNAP-depleted mouse ESCs (Jiang et al, 2020). They are also in line with observations of transcriptional inhibitors weakening, but not abolishing, TAD boundaries (Hug et al, 2017; Barutcu et al, 2019). Second, how does RNAP depletion promote formation of hundreds of *de novo* loops that are also longer? According to our simulations, and pending on the direction of elongation, RNAPs can reel DNA such that it counters extrusion while also acting as physical blockades to it. This is reminiscent of the condensin-polymerase antagonism reported for bacteria and flies (Brandão et al, 2019; Rowley et al, 2019), and recently observed using super-resolution imaging in mESCs (Gu et al, 2020), offering orthogonal validation to our Hi-C data. Some of these newly emerging loops form on the basis of strengthened Polycomb interactions (**Figure S5A,B**), justified by the increase in H3K27me3 levels and in line with recent work, where cohesin removal had the same result (Rhodes et al, 2020). As regards increased loop lengths, our simulations argue that this is a consequence of reduced cohesin loading rates to chromatin, as well as of the different loading patterns in wild-type versus RNAP-depleted cells. These translate into fewer cohesin rings acting on the fiber at any given time, and such reduced “crowding” creates fewer extrusion conflicts and allows longer loops to form (with reduced unloading also promoting loop enlargement; Haarhuis et al, 2017; Gassler et al, 2017). Finally, why are the effects of RNAPII depletion more obvious upon G1 reentry? We believe that this is due to a combination of effects. On one hand, cell synchronization counters the inherent heterogeneity of contacts in individual cells. On the other, early chromatin refolding and transcription bursts in the mitosis-to-G1 transition suggest that RNAPs preempt a central role in establishing a loop-based chromosomal architecture by instructing cohesin loading and setting up compartments.

In summary, we uncovered a dependency of loop extrusion on RNAPII that predominates genome reorganization following exit from mitosis. This dependency comes to add to the significance of mitotic bookmarking, as transcription factor association with mitotic chromatin may dictate RNAP positioning and, in a next step, cohesin loading and loop extrusion. Nonetheless, the precise interplay between polymerases, transcription factors and cohesin subunits during this transition remains to de elucidated.

## AUTHOR CONTRIBUTIONS

SZ, NÜ, and NJ performed experiments. SZ, NJ, and EGG performed bioinformatics analyses. GF, MC, and DM performed computational modeling. HG and VR performed 3D DNA FISH. JS, ABH, and KSW performed super-resolution STORM imaging. CB and JA performed next-generation sequencing. AP conceived and supervised the study, and compiled the manuscript with input from all coauthors.

## ACKNOWLEDGEMENTS

We thank Peter Cook, Nick Gilbert, and Juanma Vaquerizas for critical reading of this manuscript, and the NGS Integrative Genomics core unit of the UMG for sequencing. Work in the laboratory of AP is funded via the Deutsche Forschungsgemeinschaft (DFG) via the SPP2202 (PA2456/11-1) and TRR81 programs (INST160/697-1). SZ is supported by a CSC doctoral fellowship. SZ, NÜ, and NJ are further supported by the International Max Planck Research School for Genome Science. Work in the lab of DM is supported by an ERC Consolidator grant, and in that of VR by the DFG via the SFB1361 (393547839) and SPP2202 programs (402733153).

## METHODS

### Cell synchronization and sorting

mAID-POLR2A(RPB1)-mClover DLD-1 (previously described; Nagashima et al, 2019) and TO2B^-/-^-TOP2A-mAID HCT116 cells (Gothe et al, 2019) were passaged in RPMI-1640 medium supplemented with 10% FBS under 5% CO2. Inducible depletion of RPB1 or TOP2A initiated via treatment with doxycycline for 24 h to induce *TIR1* expression, before addition of 500 μM indole-3-acetic acid solution (“auxin”, Sigma-Aldrich) for different times to induce RPB1 degradation. For cell synchronization, G2/M arrest was achieved by addition of 10 μM RO-3306 inhibitor for 21 h. Following this incubation time, cells were washed with PBS, and auxin-supplemented medium was added for up to 6 h to allow cells to quantitatively enter G1. At this point, synchronized or asynchronous cells treated with auxin for up to 14 h were harvested, where applicable resuspended in 1 μg/ml propidium iodide to counterstain DNA, and sorted to isolate G1 cells using a FACS Canto II flow cytometer (Becton Dickinson).

### In situ Hi-C and data analysis

All *in situ* Hi-C was performed using the Hi-C+ kit (Arima Genomics) as per manufacturer’s instructions. The resulting Hi-C libraries were paired-end sequenced on a NovaSeq6000 platform (Illumina) to >500 million read pairs per replicate (**Table S1**). Reads were separately aligned to the reference build of the human genome (hg38) using BWA via the Juicer (v. 1.11.09) software to generate .hic files (Durand et al, 2016). Only reads with high MAPQ (>30) were considered for further analysis, and bin-to-bin interactions were extracted from KR-balanced matrices using the “dump” utility of Juicer at different resolutions from .hic files. A-/B-compartment stratification was performed using the “eigenvector” utility of Juicer on 250 kbp-resolution matrices, and both gene and H3K27ac ChIP-seq signal density (Abraham et al, 2017) were used to deduce A-compartments, before assigning positive scores to them. Saddle plots were generated as described previously (Zhang et al, 2019). For topologically-associating domains (TADs), KR-balanced matrices we processed via a combination of “directionality index” plus HMM tools at 10 kbp-resolution and in 500-kbp windows as described previously (Rao et al, 2014). In a last step, TADs smaller than 150 kbp or those in centromeric regions were filtered out. Insulation scores at TAD boundaries were calculated using a sliding 120 kbp x 120 kbp window along the matrix diagonal at 10-kbp resolution as previously described (Crane et al, 2015); squares with a sum of interactions <12 were filtered out. For loop detection, we used SIP (Rowley et al, 2020) with the following parameters: *-res 10000 -mat 2000 -g 2 -d 3 -fdr 0.01 -nbZero 4 -cpu 1 -factor 1 -max 2 -min 2 -sat 0.01 -t 2800 -norm KR -del true*, and an FDR <0.01 to stringently filter the resulting loop lists. Loops specific to a given condition were determined using *pgltools* (with *-d* 29999 due to the smallest allowable loop size of 30 kbp; Greenwald et al, 2017). Finally, aggregate peak plots were generated using the APA utility in Juicer to calculate with standard parameters (*-r 10000 -k KR -q 3 -w 6 -n 15 -u*), before scaling from 0-2 to facilitate comparison. All code is available at: https://github.com/shuzhangcourage/HiC-data-analysis.

### High throughput 3D-DNA fluorescence in situ hybridization (FISH)

Dual color DNA FISH was performed using the BAC probes targeting different chromosomes (**Table S2**) and labeled with Alexa488-dUTP, Alexa568-dUTP or Alexa647-dUTP by nick translation on G1-sorted control and auxin-treated DLD-1 reentry cells seeded on glass slides. Images were acquired using an Opera Phenix High Content Screening System (PerkinElmer), equipped with four laser lines (405 nm, 488 nm, 568 nm, and 640 nm) and two 16-bit CMOS cameras. Images for 3D and radial distances were acquired in confocal mode using a 40X water objective (NA 1.1), and analyzed as described previously (Zirkel et al, 2018). DNA content was also analyzed as previously described (Roukos et al, 2015).

### Chromatin immunoprecipitation (ChIP) coupled to sequencing

DLD-1 cells cultured to 80% confluence in 15-cm dishes were crosslinked in 1% PFA/PBS 10 min at room temperature. Cells were processed using the NEXSON ChIP protocol as previously described (Arrigoni et al, 2015). In brief, nuclei were isolated via sonication using a Bioruptor Pico (Diagenode; 9 cycles of 10 sec *on* and 30 sec *off*). Chromatin was then sheared in the recommended shearing buffer (27-30 cycles, 30 sec *on* and 30 sec *off*) to a range of 200-500 bp-long fragments, and immunoprecipitation was performed using 4 μg of the appropriate antibody (anti-CTCF: 61311, Active Motif; anti-RAD21: ab88572, Abcam; anti-GFP: ab290, Abcam). Paired-end sequencing was performed on a NovaSeq6000 platform (Illumina) yielding >25 million reads per sample. Raw reads were processed using the pipeline by the ENCODE Data Coordinating Center (DCC; v. 1.5.0, https://github.com/ENCODE-DCC). Briefly, reads were aligned to the hg38 human genome build using Bowtie2 (Langmead and Salzberg, 2012) and filtered with SAMtools (Li et al, 2009) to remove unmapped, non-primary and duplicate reads. Peak calling was performed via SPP (Kharchenko et al, 2008) and IDR (Li et al, 2011) was used to threshold peak lists. Input-normalized signal tracks were generated using MACS2 (Zhang et al, 2008) and coverage plots and heatmaps using Deeptools (Ramírez et al, 2014).

### Assay for Transposase-Accessible Chromatin using sequencing (ATAC-seq) and data analysis

Tn5 transposase-accessible chromatin was isolated from human DLD-1 mAID-RPB1 cells according to the standard ATAC-seq protocol (Buenrostro et al, 2015) with one modification aiming at quantitative scaling of the resulting data. In brief, 10^5^ DLD-1 cells per replicate we “spiked” with 200 *D. melanogaster* S2 cells, washed in 1x PBS and added to lysis buffer (10 mM Tris-HCl pH 7.4, 10 mM NaCl, 3 mM MgCl_2_, 0.1% NP-40, 0.1% Tween-20, and 0.01% digitonin) for 3 min (Corces et al, 2017) to isolate nuclei. Nulcei were next washed in washing buffer (10 mM Tris-HCl pH 7.4, 10 mM NaCl, 3 mM MgCl_2_, 0.1% Tween-20) and pelleted by centrifugation. Isolated nuclei were resuspended in transposase reaction mix (25 μl 2x TD buffer, 16.5 μl 1x PBS, 0.5 μl 10% Tween-20, 0.5 μl 1% digitonin, 2.5 μl Tn5, and 5 μl nuclease-free H_2_O) and incubated at 37°C for 30 min on Thermomixer under constant shaking at 1000 rpm. The transposition reaction was terminated by the addition of stop buffer (50 mM Tris-HCl pH 8, 10 mM EDTA, 1% SDS), and DNA purified using the DNA Clean & Concentrator kit (Zymo Research). Following standard library generation, samples were sequenced to >40 million read pairs on a NovaSeq6000 platform (Illumina). Read pairs were mapped to the hg38 and dm6 reference genome builds for human and Drosophila, respectively, using Bowtie2 (Langmead and Salzberg, 2012). Non-primary, unmapped, duplicate, and mitochondrial reads were removed. Data mapping to the Drosophila genome were used in ChIPseqSpike (https://bioconductor.org/packages/release/bioc/html/ChIPSeqSpike.html) for calculating scaling factors in order to produce RPKM-normalized and scaled coverage files.

### Immunofluorescence and image quantification

DLD-1 cells grown on coverslips were fixed in 4% PFA/PBS for 10 minutes at room temperature. After washing once in PBS, cells were permeabilized with 0.5% Triton-X/PBS for 5 min at room temperature, washed three times in PBS, blocked using 1% BSA for 1 h, and incubated with the appropriate primary antibody for 1 h at room temperature (anti-RNAPII: 1:500, 61086, Active Motif; anti-H3K27ac: 1:500, 39133, Active Motif; anti-H3K27me3: 1:500, 39155, Active Motif; anti-RAD21: 1:800, ab992, Abcam; anti-Fibrillin: 1:100, sc-393968, Santa Cruz). For visualizing nascent transcripts, cells were pre-incubated with 3 mM 5-ethynyl uridine (EU) for 30 min at 37°C in their growth medium, fixed and processed with the Click-iT EdU chemistry kit (Invitrogen). Images were acquired on an Leica dmi8 microscope using the LASX software. Quantification of nuclear fluorescence were performed by drawing a mask based on DAPI staining, and then calculating the mean intensity per area falling under this mask. Colocalization was assessed using the ImageJ plugin, JACoP.

### Chromatin fractionation and western blotting

For assessing protein abundance in different sample preparations, approx. 10^6^ cells were gently scraped off 15-cm dishes, and pelleted for 5 min at 600 x *g* at room temperature, supernatants were discarded, and pellets resuspended in 100 μl of ice-cold RIPA lysis buffer (20 mM Tris-HCl pH 7.5, 150 mM NaCl, 1 mM EDTA pH 8.0, 1 mM EGTA pH 8.0, 1% NP-40, 1% sodium deoxycholate) containing 1x protease inhibitor cocktail (Roche). Next, lysates were incubated for 20 min on ice and centrifuged for 15 min at >20,000 x *g* to pellet cell debris to collect the supernatants. The concentration of each protein extract was determined using the Pierce BCA Protein Assay Kit (Thermo Fisher Scientific). For fractionation, the protocol previously described was used (Watrin et al, 2006). Following protein separation on precast SDS-PAGE gels (BioRad), proteins were detected using different primary antibodies (see **Table S3**), and signal was detected with the Pierce SuperSignal West Pico ECL kit (Thermo Fisher Scientific).

### Factory RNA sequencing and data analysis

Nascent RNA from ^~^10 million mAID-RPB1-mCLover DLD-1 or TO2B^-/-^-TOP2A-mAID HCT116 cells was isolated according to “factory-seq” protocol (Melnik et al, 2016). Briefly, cells were gently scraped and lysed in isotonic “physiological buffer” supplemented with 0.5% NP40 buffer. After assessing lysis and nuclei integrity on a hemocytometer microscopy, nuclei were treated with DNase I (Worthington) for 30 min at 33°C, washed, and nuclei were lysed in “native lysis buffer” and treated with caspase group III enzyme mix (PromoKine), pelleted by centrifugation, before the supernatant holding nascent RNA was collected in TRIzol (Invitrogen) and purified using the Direct-Zol RNA purification kit (Zymo). Following standard strand-specific cDNA library preparation using the TruSeq kit (Illumina), sequencing was performed on a NovaSeq6000 platform (Illumina) to >40 million paired-end reads. Raw reads were mapped to human genome (build hg38) using STAR (Dobin et al, 2013), quantified using *FeatureCounts* (Liao et al, 2014) considering only primary aligned, properly paired reads, and normalized using RUVseq (Risso et al, 2014). Differential gene expression was computed using DESeq2 (Love et al, 2014). For gene set enrichment, GSEA (Subramanian et al, 2005) was ran on significantly changing genes (*P*_adj_≤0.05) that are listed in **Table S4**.

### Dual-color super-resolution dSTORM imaging and analysis

DLD-1 control and auxin-treated reentry cells were seeded onto coverslips, fixed, stained, and imaged as described previously (Casa et al, 2020). In brief, fixed and immunostained cells for NIPBL and CTCF (as described above) were mounted to an Attofluor cell chamber (Thermo Fisher Scientific) and in 1 mL of *d*STORM buffer (25 mM MEA, glucose oxidase, 50 mM NaCl, and 10% glucose in 10 mM Tris-HCl pH 8.0). The chamber is then sealed with a coverslip and left on the microscope at room temperature for 30 min prior to imaging, to minimize drift. Imaging was performed on a Zeiss Elyra PS1 system fitted with an Andor iXon DU 897, 512 × 512 EMCCD camera. Images were made using a 100× 1.49NA TIRF objective in HiLo mode. Movies of 12,000 frames were recorded with an exposure time of 33 msec. Multichannel images were acquired sequentially from high wavelength to lower wavelengths. *d*STORM movies for each protein target were analyzed via the Zeiss ZEN 2012 software, and localizations with a precision of >50 nm were discarded. All remaining localizations were drift-corrected using a model-based approach. All additional analysis was done in R (https://www.R-project.org/), localizations from individual nuclei were clustered based on their density using a kernel density estimation (KDE)-based clustering algorithm with the threshold set to 0.05 for all channels. The areas of CTCF or NIPBL clusters were measured using the KDE binary image, and distances between closest neighbors calculated.

### Computational modeling

For the 3D simulations, we used our previously described HiP-HoP model (Buckle et al, 2018), extended to account for interactions in inactive regions. This model combines our initial “transcription factory” model (Brackley et al, 2016) with loop extrusion (Fudenberg et al, 2016), while also accounting for the heteromorphic nature of the chromatin fiber, which means that the local compaction (in DNA base pairs per nanometer) varies along the fiber. Here, we modeled a 10-Mbp region of HUVEC chr14 as a bead-spring polymer containing *N* beads of diameter *σ*, each representing 1 kbp of chromatin. We allowed beads in the polymer to interact via three potentials: (i) a Weeks-Chandler-Andersen (WCA) potential, which provides excluded volume interactions; (ii) a finitely-extensible-nonlinear-elastic (FENE) potential accounting for chain connectivity; and (iii) a Kratky-Porod potential describing the flexibility of the chain with parameters set to give a persistence length of 4-5 kbp (in line with that of chromatin *in vivo*). To model the heteromorphic nature of the chromatin fiber in a simple way, we included additional springs (with constants of 200 *kBT/σ^2^*) between next-to-neighbor chromatin beads along the chain (i.e., bead *i* and *i*+2) which are not associated with H3K27ac marks. As H3K27ac marks correlate with active euchromatin regions, these springs cause a local crumpling of the polymer in inactive chromatin fragments, or equivalently a swelling in active regions accounting for their generally more open conformation. Transcription factors (TFs) were simulated as diffusing beads interacting with each other via steric repulsion, again modeled via WCA potentials. We considered three types of TFs: (i) generic active TFs bind strongly (potential depth 7.9 *k_B_T*) to chromatin beads associated with accessible chromatin (defined using ENCODE DHS-seq data), and weakly to beads associated with H3K27ac (potential depth 3.4 *k_B_T*); (ii) HP1-like inactive TFs bind to beads enriched in H3K9me3 marks (potential depth 3.4 *k_B_T*); and (iii) Polycomb-like TFs bind to beads enriched in H3K27me3 marks (potential depth 7.9 *k_B_T*). All of these interactions were modeled via a truncated-and-shifted Lennard-Jones (LJ) potential. To account for post-translational modifications, we allowed each TF type to switch between a binding and a non-binding state at a defined rate (*k*_sw_ = 10^−3^*τ*^−1^, where *τ* is the simulation time unit). The binding state was characterized by the aforementioned interaction strengths, whereas the non-binding state only by steric interactions with chromatin beads (via WCA potentials). We considered 250 active TFs, 625 HP1-like TFs, and 125 Polycomb-like TFs in the wild-type simulations, whereas the RNAPII-degron simulations were run without active TFs. We also implemented non-specific interactions between inactive chromatin beads (via a truncated-and-shifted LJ potential with depth 0.45 *k_B_T*) to account for the generic “phase separation” between eu- and hetero-chromatin. Finally, loop extrusion was modeled by representing cohesin dimers as further additional springs. Loop extrusion dynamics were determined by the number density of cohesins (*n_c_* = 0.01/kbp) on the chromatin fiber and two rates: the unbinding rate (*k*_off_ = 2.5×10^−5^*τ*^−1^) and the extrusion rate (*v* = 4×10^−3^ kbp/*τ*^−1^). Upon binding of cohesin, we introduced an additional spring between two nearby beads along the fiber (*i* and *i*+3, since crumpling springs already link *i* and *i*+2); the equilibrium length and the spring constant of cohesin bonds were set to 1.5 *σ* and 40 *kBT/σ^2^*, respectively. When cohesins were removed from chromatin, they were instantly repositioned along the fiber. Wild-type conditions were simulated by background random loading (with 10% probability) but with predominant loading at DHS beads (with 90% probability); the RNAPII-degron was simulated by only considering random loading. Finally, a cohesin halted either upon colliding with another extruding complex or upon reaching a CTCF site whose direction was against the direction of extrusion (as seen experimentally; Rao et al, 2014). Note that CTCF sites and orientation were obtained by ENCODE tracks, taking care to include in our simulations only sequences overlapping cohesin (RAD21) peaks – this procedure singles those CTCF binding sites that are relevant to looping. All constituents of the system (chromatin beads and TFs) were allowed to diffuse, and their dynamics were governed by a Langevin equation as described before (Buckle et al, 2018), and implemented using Python and the LAMMPS molecular dynamics software package (Plimpton, 1995) as a library.

For the 1D simulations, we considered a 3 Mbp-long chromatin fiber coarse-grained into segments of 1 kbp. Again, we modeled data from a specific subregion of HUVEC chr14. We simulated the dynamics of cohesin rings (total number *N_rings_* = 30), each of which could be in one of two states: either bound (i.e., on the fiber) or unbound (i.e., in the diffuse pool). Binding and unbinding were modeled as stochastic processes with rates *k*_on_ and *k*_off_, respectively. When on the fiber, a cohesin molecule was modeled as a dimer, with each monomer undergoing active extrusion at speed *v*. Each monomer could proceed until it hit a CTCF site with orientation conflicting with its direction of travel, at which point it became immobile (Fudenberg et al, 2016). If a cohesin complex was halted on one side, its other side could continue to move independently. When both monomers in a cohesin dimer became stuck at convergent CTCF sites, the unbinding rate of the dimer was decreased by a factor of 10 to model CTCF-mediated stabilization of extruded chromatin loops. We let monomers in a cohesin dimer interact with each other via steric exclusion so that extrusion would be halted temporarily if another monomer was in their way. Wild-type conditions were simulated by assuming that cohesin was loaded as described above, with 10% background random loading and 90% at DHS sites, whereas RNAPII-degron conditions were simulated by only retaining the random background loading. To simulate feedback of transcription on extrusion, we assumed that the speed of extrusion was reduced by a factor *f* when the direction of extrusion and that of transcriptional elongation of an active gene were conflicting. Parameters in the simulations were set to *k*_on_ = 2×10^−2^, *k*_off_ = 10^−3^, *k*_on_ = 2×10^−2^, *k*_off_ = 10^−3^, *v* = 0.16 for the wild-type, and to *k*_on_ = 2×10^−3^, *k*_off_ = 10^−3^, *v* = 0.16 for the RNAPII-degron. These values can be mapped to *k*_on_ = 1 min^−1^, *k*_off_ = 0.05 min^−1^, *v* = 0.133 kbp/s (wild-type, corresponding to a residence time on chromatin ^~^20 min) or *k*_on_ =0.067 min^−1^, *k*_off_ = 0.033 min^−1^, *v* = 0.089 kbp/s (RNAPII-degron, corresponding to a residence time on chromatin ^~^30 min). With these parameters the chromatin length explored during an extrusion event, *λ* = *v*/*k*_off_, was the same for wild-type and degron, and comparable to that used previously (Fudenberg et al, 2016). For simulations including the feedback of transcription on extrusion, we varied *f* (which is unknown experimentally) between 0.1 and 0.9 to simulate a variety of scenarios, and presented the case with *f=0.1*, which lead to the most pronounced effects.

### Statistical analyses

*P*-values associated with the Student’s t-tests and Fischer’s exact tests were calculated using GraphPad (http://graphpad.com/), those associated with the Wilcoxon-Mann-Whitney test using the EDISON-WMW tool (Marx et al, 2016). Unless otherwise stated, P-values <0.01 were deemed significant.

### Data availability

NGS data generated in this study are available via the NCBI Gene Expression Omnibus (GEO) repository.

---

**Figure S1.**
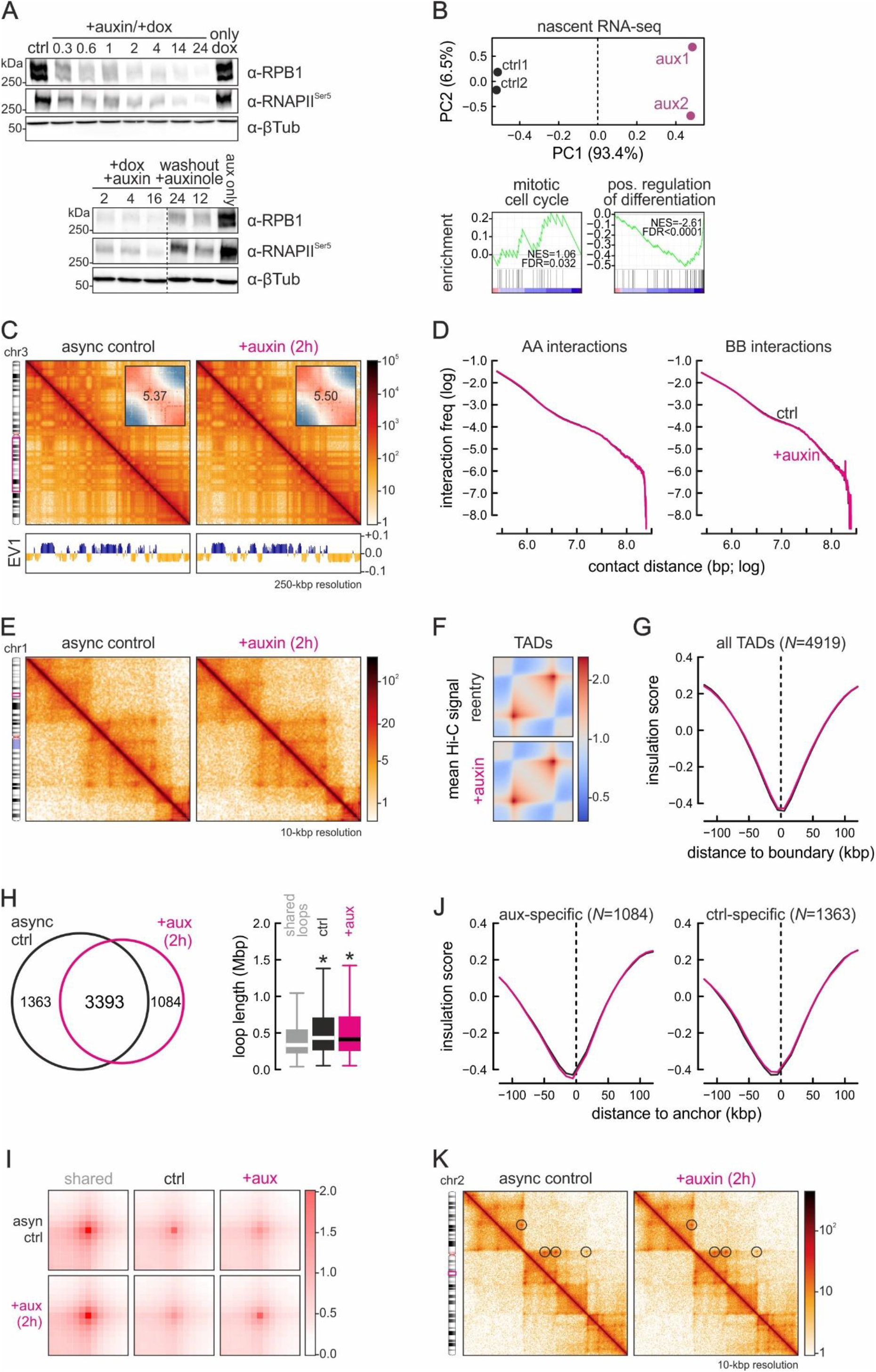
RNAPII degradation, recovery, and its effects on interphase chromatin folding. (**A**) Western blots showing depletion of total cell RPB1 or phospho-Ser5-RNAPII on increasing exposure to doxycycline plus auxin (top) or RNAPII recovery following auxin washout in the presence of auxinole (bottom); β-tubulin provides a loading control. (**B**) *Top*: PCA plot for G1-sorted control (black) and 14-h auxin-treated nascent RNA-seq data (purple). *Bottom*: Gene set enrichment analysis. (**C**) Exemplary Hi-C maps of a subregion of chr3 from asynchronous control (left) and 2-h auxin-treated DLD1-mAID-RPB1 cells (right) at 250-kbp resolution aligned to first eigenvector values (below). *Insets*: saddle plots showing no change in A/B-compartment insulation. (**D**) Decay plots showing Hi-C interaction frequency between A- (left) or B-compartments (right) as a function of genomic distance (log) in control (black line) or 2-h auxin-treated cells (magenta line). (**E**) Exemplary Hi-C maps of a subregion of chr1 from control (left) and auxin-treated cells (right) at 10-kbp resolution. (**F**) Heatmaps showing mean TAD-level interactions in control (top) and auxin-treated cells (bottom). (**G**) Line plots showing mean insulation score from control (black line) and auxin-treated cells (magenta line) in the 240 kbp around all TAD boundaries in control cells. The number of TAD boundaries queried (*N*) is indicated. (**H**) *Left*: Venn diagram showing shared and unique loops in control (black) and auxin-treated Hi-C data (magenta). *Right*: Loop lengths displayed as boxplots. *: significantly different; *P*<0.01, Wilcoxon-Mann-Whitney test. (**I**) APA plots showing mean Hi-C signal for shared (left), control- (middle), and degron-specific loops (right) from panel H. (**J**) As in panel G, but for the anchors of control- (left) and degron-specific loops (right; from panels H,I). **(K)** Exemplary Hi-C maps showing little change in loop emergence (circles) between control (left) and auxin-treated cells (right).

**Figure S2.**
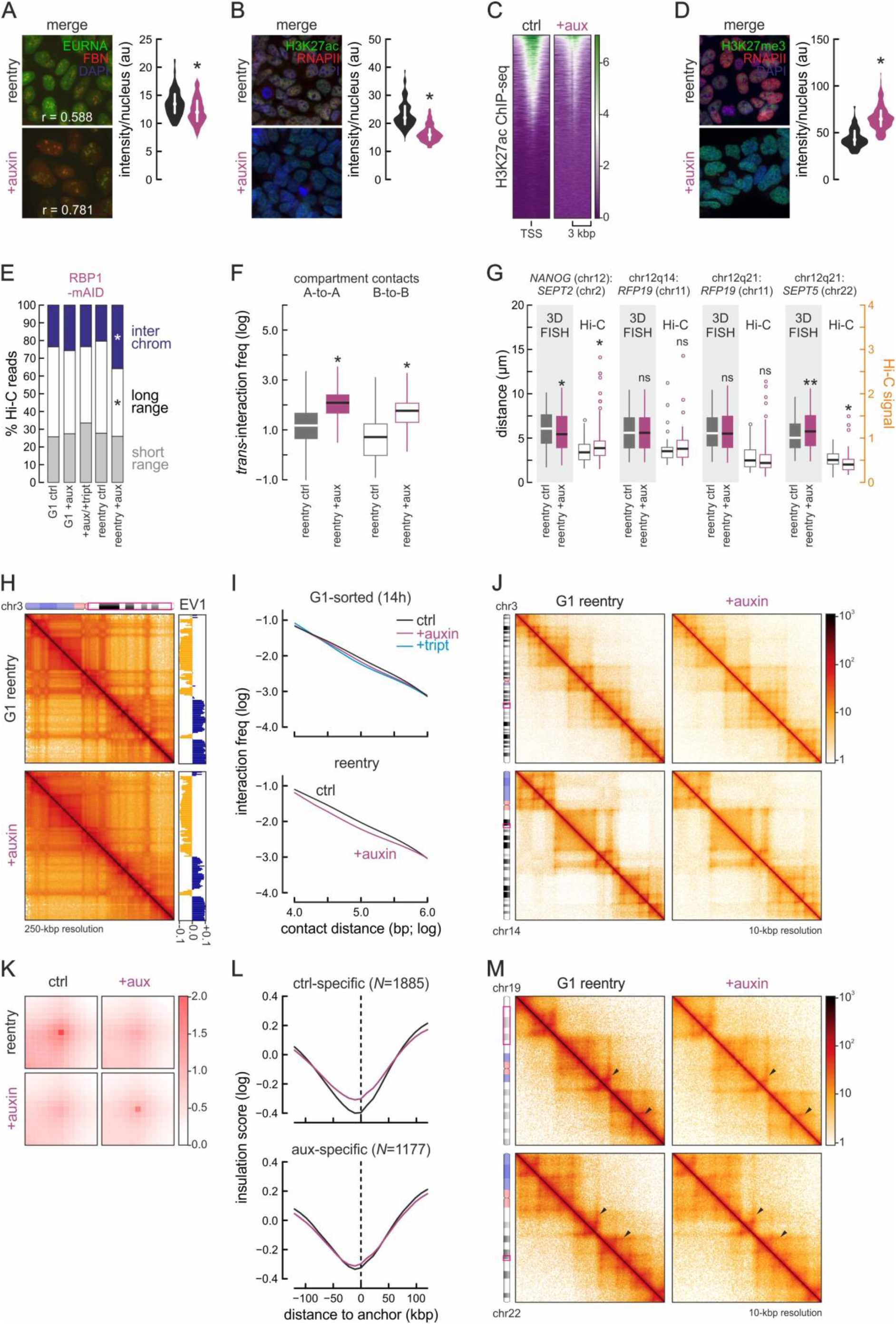
RNAPII degradation affects chromatin refolding in *cis* and in *trans* following mitotic exit. (**A**) *Left*: Exemplary widefield immunofluorescence images of DLD1-mAID-RPB1 G1-reentry cells treated with doxycycline plus auxin (bottom) or not (top) and stained for fibrillin and EU-labeled nascent RNA; nuclei were counterstained with DAPI. *Right*: Bean plots showing EU-RNA fluorescence per nucleus. *: significantly different; *P*<0.01, Wilcoxon-Mann-Whitney test. (**B**) As in panel A, but stained for RNAPII and H3K27ac, and quantifying H3K27ac levels. (**C**) Heatmaps showing H3K27ac ChIP-seq signal in the 6 kbp around active TSSs in control versus auxin-treated reentry cells. (**D**) As in panel A, but stained for RNAPII and H3K27me3, and quantifying H3K27me3 levels. (**E**) Bar plots showing the percent of Hi-C reads representing interchromosomal (blue), long-range (>20 kbp) or short-range intrachromosomal contacts (white) across Hi-C datasets. *: significantly different; *P*<0.01, Fischer’s exact test. (**F**) Boxplots showing interchromosomal interactions between A-A and B-B compartments in auxin-treated versus control reentry cells. *: significantly different; *P*<0.01, Wilcoxon-Mann-Whitney test. (**G**) Boxplots comparing changes in interchromosomal distances for the loci indicated assessed using high throughput 3D-DNA FISH (grey background) and Hi-C data at 0.5-Mbp resolution. *: significantly different; *P*<0.01, Wilcoxon-Mann-Whitney test. (**H**) Additional Hi-C examples of a subregion of chr3 from control (top) and auxin-treated reentry cells (bottom) at 250-kbp resolution aligned first eigenvector values (right). (**I**) Decay plots showing Hi-C interaction frequency as a function of genomic distance (log) at the scale of TADs (0.01-1 Mbp) in control (black line) or auxin-treated reentry cells (purple/blue lines). (**J**) Additional Hi-C examples of subregions in chr3 and 14 from control (left) and auxin-treated reentry cells (right) at 10-kbp resolution. (**K**) APA plots showing mean Hi-C signal for loops lost/gained in control (left) and auxin-treated reentry cells (right). (**L**) Line plots showing mean insulation scores in the 240 kbp around control- (top) or degron-specific loops (bottom) from control (black line) and auxin-treated cells (purple line). The number of anchors queried (*N*) is indicated. (**M**) Hi-C maps showing exemplary changes in “stripes” (arrowheads) between control (left) and auxin-treated reentry cells (right) at 10-kbp resolution.

**Figure S3.**
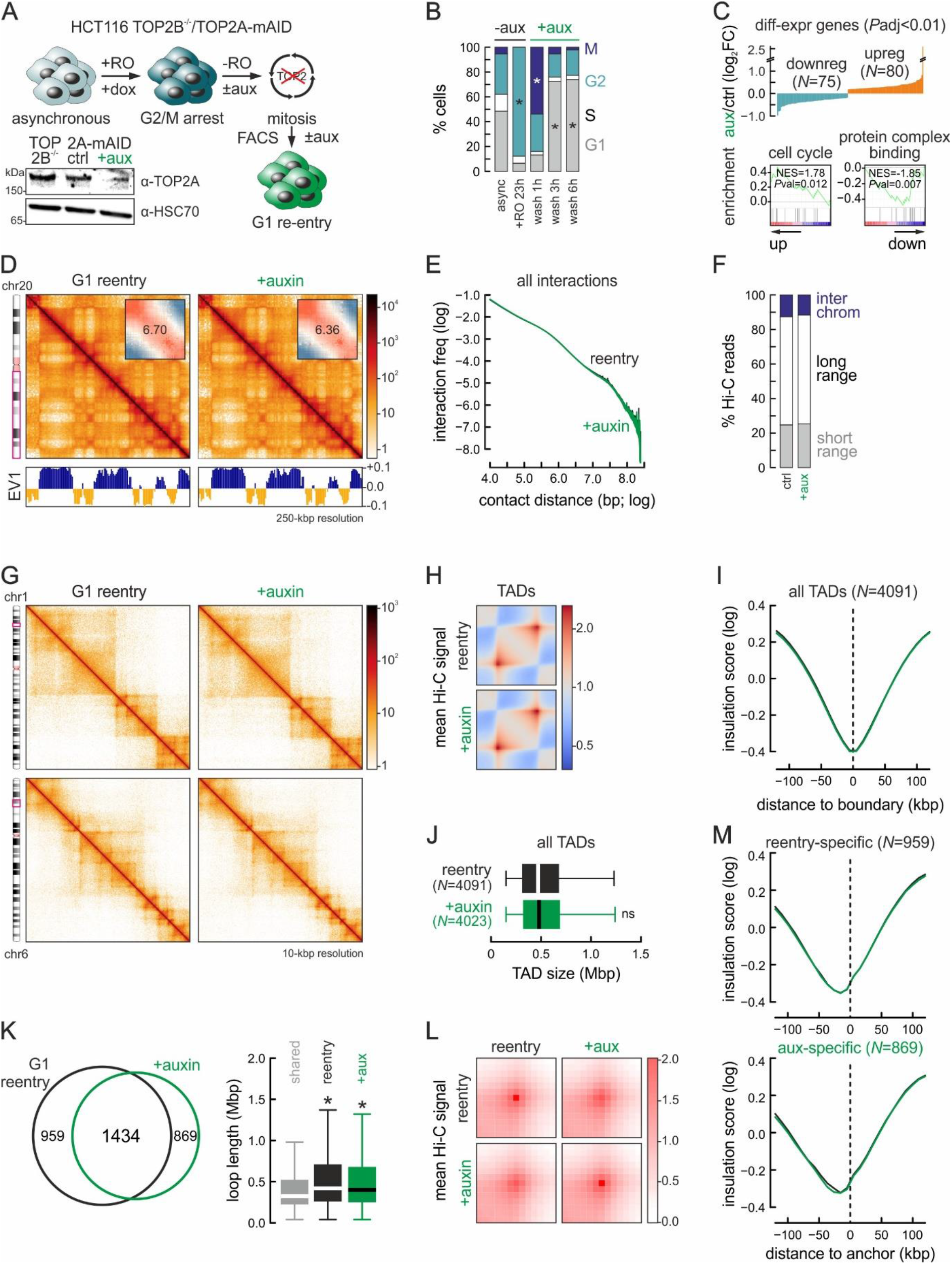
Topoisomerase II depletion marginally affects chromatin refolding following mitotic exit. (**A**) *Top*: Overview of the experimental scheme for HCT116-TOP2B^-/-^-TOP2A-mAID cell synchronization and release. *Bottom left*: Western blots showing auxin-mediated TOP2A degradation; HSC70 provides a loading control. (**B**) Bar plots showing the percent of cells in each cell cycle phase from panel A. *: significantly different; *P*<0.01, Fischer’s exact test. (**C**) *Top*: Graph showing nascent transcription changes (log_2_ fold-change compared to control cells, *P*_adj_<0.01) upon TOP2B-degradation. *Bottom*: Gene set enrichment analysis. (**D**) Exemplary Hi-C maps of a subregion of chr20 from control (left) and auxin-treated G1-reentry cells (right) at 250-kbp resolution aligned to first eigenvector values (below). *Insets*: saddle plots showing A/B-compartment insulation. (**E**) Decay plots showing Hi-C interaction frequency between compartments as a function of genomic distance (log) in control (black line) or auxin-treated reentry cells (green line). (**F**) Bar plots showing the percent of Hi-C reads in interchromosomal (blue) or long- (>20 kbp) and short-range intrachromosomal contacts (white) in control and auxin-treated reentry Hi-C data. *: significantly different; *P*<0.01, Fischer’s exact test. (**G**) Exemplary Hi-C maps of subregions in chr1 and 6 from control (left) and auxin-treated reentry cells (right) at 10-kbp resolution. (**H**) Heatmaps showing mean TAD-level interactions in control (top) and auxin-treated reentry cells (bottom). (**I**) Line plots showing mean insulation scores from control (black line) and auxin-treated reentry cells (green line) in the 240 kbp around all TAD boundaries. The number of TADs queried (*N*) is indicated. (**J**) Boxplots showing TAD size changes between control (black) and auxin-treated reentry cells (green). (**K**) *Left*: Venn diagram showing shared and unique loops between control (black) and auxin-treated reentry cells. *Right*: Loop lengths displayed as boxplots. *: significantly different; *P*<0.01, Wilcoxon-Mann-Whitney test. (**L**) APA plots showing mean Hi-C signal for loops lost/gained in control (left) and auxin-treated reentry cells (right), from panel K. **(M)** As in panel I, but for loop anchors from panels K,L.

**Figure S4.**
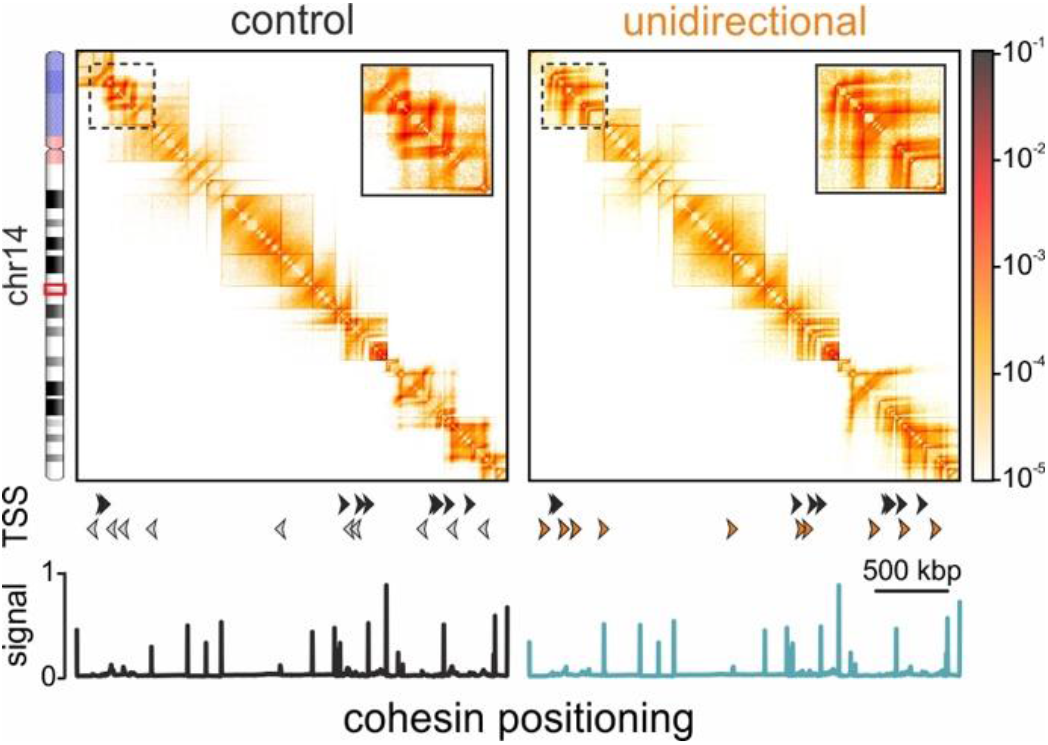
The direction of transcription can affect loop extrusion. Heatmaps rendered from loop extrusion 1D simulations representing wild-type cells (left) or cells where all TSS are transcribed in the same direction (right) in the HUVEC chr14:53-56 Mbp segment. Profiles of cohesin positioning and TSS orientations are aligned below each heatmap (reoriented TSSs are indicated in orange).

**Figure S5.**
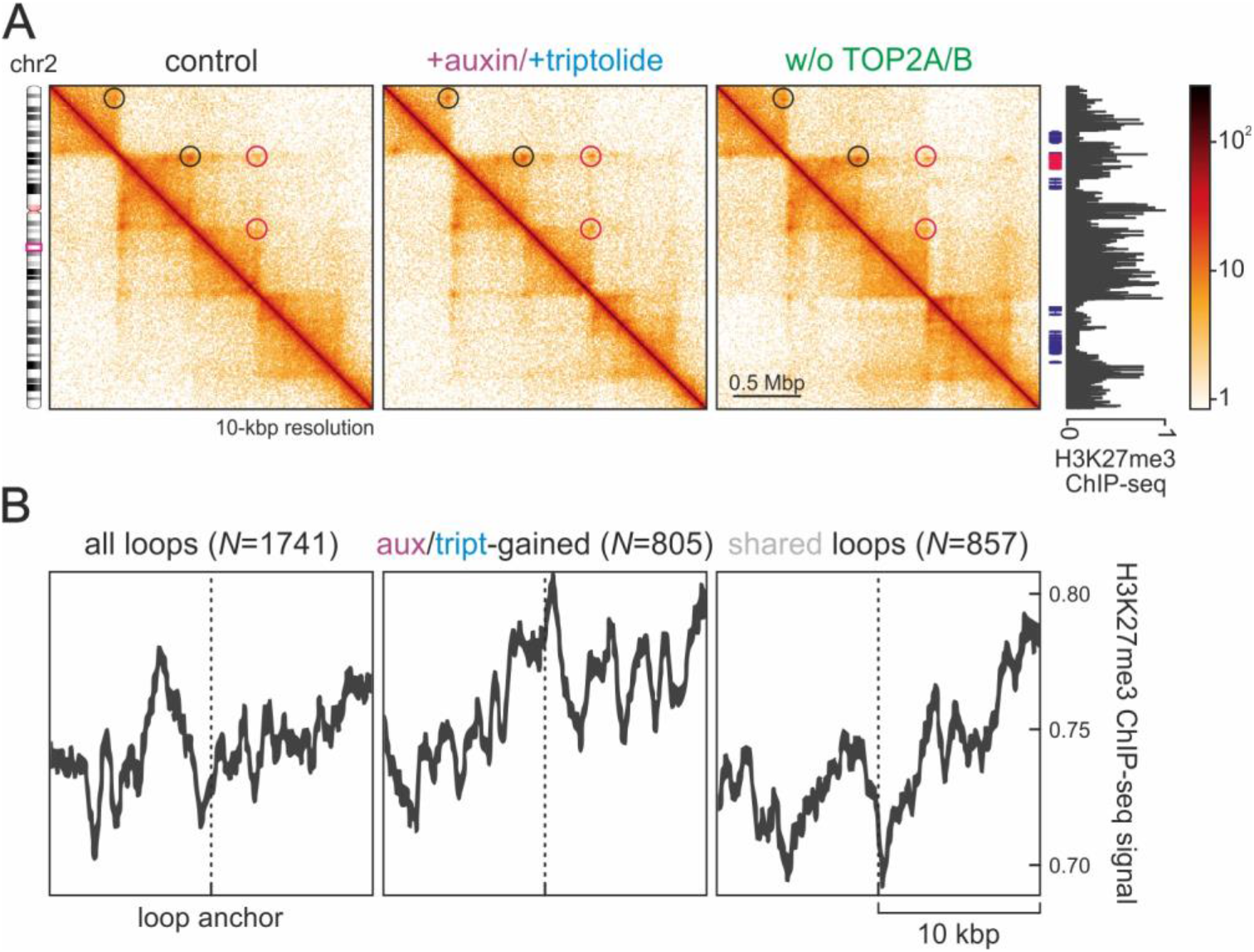
RNAPII depletion accentuates H3K27me3-anchored interactions. (**A**) Hi-C maps from control (left), auxin-/triptolide-treated (middle) or TOP2A/B-depleted cells (right) in the chr2 subregion encompassing the *HOXD* gene cluster. H3K27me3 ChIP-data from control cells are aligned to the maps, and H3K27me3-anchored loops emerging are denoted (red circles). (**B**) Line plots showing mean H3K27me3 ChIP-seq signal in the 20 kbp around all loops (left), auxin- /triptolide-gained loops (middle) or loops shared between RNAPII-depleted and control cells (right). The number of loops in each group (*N*) is indicated.

## SUPPLEMENTAL TABLES

**Table S1.** General statistics of all Hi-C datasets.

| Hi-C dataset: | total reads | % aligned | Hi-C contacts | % inter-chromo | % intra-chromo | % short-range | % long-range |
| --- | --- | --- | --- | --- | --- | --- | --- |
| 2h control | 570,638,421 | 89.91 | 387,244,720 | 10.44 | 57.42 | 21.94 | 35.47 |
| 2h +auxin | 521,206,158 | 91.46 | 361,994,319 | 10.84 | 58.62 | 22.43 | 36.17 |
| 14h control | 512,551,503 | 89.70 | 339,693,791 | 16.49 | 49.78 | 16.75 | 33.03 |
| 14h +auxin | 516,427,195 | 89.04 | 338,221,913 | 17.69 | 47.80 | 17.65 | 30.14 |
| 14h +aux/+tript | 461,569,471 | 90.67 | 313,489,486 | 16.87 | 51.04 | 19.88 | 28.67 |
| reentry control | 1,129,688,081 | 89.57 | 740,115,313 | 14.26 | 51.26 | 17.86 | 33.39 |
| reentry +auxin | 1,691,677,000 | 89.59 | 1,069,866,689 | 23.41 | 39.83 | 16.14 | 23.69 |
| TOP2 control | 541,077,797 | 88.39 | 364,665,469 | 10.32 | 57.07 | 23.96 | 33.11 |
| TOP2 +auxin | 546,798,121 | 88.24 | 368,526,565 | 10.08 | 57.32 | 24.49 | 32.83 |

**Table S2.** Targets of 3D-DNA FISH probes.

| probe target: | genomic coordinates (hg19) |
| --- | --- |
| *Nanog* | chr12:7,696,106-7,696,694 |
| chr12q14 | chr12:60,211,422-60,211,814 |
| chr12q21 | chr12:83,646,182-83,815,279 |
| *PRPF19* | chr11:60,677,496-60,677,988 |
| *SEPT2* | chr2:242,097,006-242,097,495 |
| *SEPT5* | chr22:19,578,750-19,579,058 |

**Table S3.** Antibodies used for Western blotting.

| antibody: | cat. No., provider | working dilution |
| --- | --- | --- |
| anti-CTCF | 61311 Active Motif | 1:2,000 |
| anti-GFP | ab290, Abcam | 1:1,000 |
| anti-H3K27ac | 39133, Active Motif | 1:1,000 |
| anti-H3K27me3 | 39155, Active Motif | 1:1,000 |
| anti-HSC70 | sc-7298, Santa Cruz | 1:2,000 |
| anti-MAU2 | ab183033, Abcam | 1:1,000 |
| anti-NIPBL | A301-779A, Bethyl | 1:10,000 |
| anti-RAD21 | ab992, Abcam | 1:1,000 |
| anti-RBP1 | ab817, Abcam | 1:500 |
| anti-RNAPI | sc-48385, Santa Cruz | 1:200 |
| anti-RNAPIII | ab88243, Abcam | 1:1,000 |
| anti-Ser5-RNAPII | 61086, Active Motif | 1:1,000 |
| anti-SMC1A | ab9262, Abcam | 1:4,000 |
| anti-WAPL | 16370-1-AP, Proteintech | 1:1,000 |
| anti-βTubulin | T0198, Sigma-Aldrich | 1:2,000 |

**Table S4.** Significantly up/downregulated genes in auxin-treated DLD-1 or HCT116 models (.xlsx file).

## REFERENCES

Abramo K, Valton AL, Venev SV, Ozadam H, Fox AN, Dekker J. (2019) A chromosome folding intermediate at the condensin-to-cohesin transition during telophase. Nat Cell Biol. 21:1393–1402.

Achar YJ, Adhil M, Choudhary R, Gilbert N, Foiani M. (2020) Negative supercoil at gene boundaries modulates gene topology. Nature 577:701–705.

Baranello L, Wojtowicz D, Cui K, et al. (2016) RNA polymerase II regulates topoisomerase 1 activity to favor efficient transcription. Cell 165:357–71.

Barutcu AR, Blencowe BJ, Rinn JL. (2019) Differential contribution of steady-state RNA and active transcription in chromatin organization. EMBO Rep. 20:e48068.

Brandão HB, Paul P, van den Berg AA, Rudner DZ, Wang X, Mirny LA. (2019) RNA polymerases as moving barriers to condensin loop extrusion. Proc Natl Acad Sci USA. 116:20489–20499.

Brackley CA, Johnson J, Kelly S, Cook PR, Marenduzzo D. (2016) Simulated binding of transcription factors to active and inactive regions folds human chromosomes into loops, rosettes and topological domains. Nucleic Acids Res. 44:3503–3512.

Brant L, Georgomanolis T, Nikolic M, et al. (2016) Exploiting native forces to capture chromosome conformation in mammalian cell nuclei. Mol Syst Biol. 12:891.

Buckle A, Brackley CA, Boyle S, Marenduzzo D, Gilbert N. (2018) Polymer simulations of heteromorphic chromatin predict the 3D folding of complex genomic loci. Mol Cell 72:786–797.

Bunch H, Lawney BP, Lin YF, et al. (2015) Transcriptional elongation requires DNA break-induced signalling. Nat Commun. 6:10191.

Busslinger GA, Stocsits RR, van der Lelij P, et al. (2017) Cohesin is positioned in mammalian genomes by transcription, CTCF and Wapl. Nature 544:503–507.

Canat A, Veillet A, Bonnet A, Therizols P. (2020) Genome anchoring to nuclear landmarks drives functional compartmentalization of the nuclear space. Brief Funct Genomics 19:101–110.

Caudron-Herger M, Cook PR, Rippe K, Papantonis A. (2015) Dissecting the nascent human transcriptome by analysing the RNA content of transcription factories. Nucleic Acids Res. 43:e95.

Casa V, Moronta Gines M, Gade Gusmao E, et al. (2020) Redundant and specific roles of cohesin STAG subunits in chromatin looping and transcriptional control. Genome Res. 30:515–527.

Chen FX, Xie P, Collings CK, et al. (2017) PAF1 regulation of promoter-proximal pause release via enhancer activation. Science 357:1294–1298.

Davidson IF, Goetz D, Zaczek MP, et al. (2016) Rapid movement and transcriptional re-localization of human cohesin on DNA. EMBO J. 35:2671–2685.

Davidson IF, Bauer B, Goetz D, Tang W, Wutz G, Peters JM. (2019) DNA loop extrusion by human cohesin. Science 366:1338–1345.

Denker A, de Laat W. (2016) The second decade of 3C technologies: detailed insights into nuclear organization. Genes Dev. 30:1357–1382.

El Khattabi L, Zhao H, Kalchschmidt J, et al. (2019) A pliable mediator acts as a functional rather than an architectural bridge between promoters and enhancers. Cell 178:1145–1158.

Fudenberg G, Imakaev M, Lu C, Goloborodko A, Abdennur N, Mirny LA. (2016) Formation of chromosomal domains by loop extrusion. Cell Rep. 15:2038–2049.

Gassler J, Brandão HB, Imakaev M, et al. (2017) A mechanism of cohesin-dependent loop extrusion organizes zygotic genome architecture. EMBO J. 36:3600–3618.

Gerlich D, Koch B, Dupeux F, Peters JM, Ellenberg J. (2006) Live-cell imaging reveals a stable cohesin-chromatin interaction after but not before DNA replication. Curr Biol. 16:1571–1578.

Giorgetti L, Lajoie BR, Carter AC, et al. (2016) Structural organization of the inactive X chromosome in the mouse. Nature 535:575–579.

Gothe HJ, Bouwman BAM, Gusmao EG, et al. (2019) Spatial chromosome folding and active transcription drive DNA fragility and formation of oncogenic MLL translocations. Mol Cell 75:267–283.

Gryder BE, Pomella S, Sayers C, et al. (2019) Histone hyperacetylation disrupts core gene regulatory architecture in rhabdomyosarcoma. Nat Genet. 51:1714–1722.

Gu B, Comerci CJ, McCarthy DG, Saurabh S, Moerner WE, Wysocka J. (2020) Opposing effects of cohesin and transcription on CTCF organization revealed by super-resolution imaging. Mol Cell doi:10.1016/j.molcel.2020.10.001.

Haarhuis JHI, van der Weide RH, Blomen VA, et al. (2017) The cohesin release factor WAPL restricts chromatin loop extension. Cell 169:693–707.

Hansen AS, Cattoglio C, Darzacq X, Tjian R. (2018) Recent evidence that TADs and chromatin loops are dynamic structures. Nucleus 9:20–32.

Hansen AS, Pustova I, Cattoglio C, Tjian R, Darzacq X. (2017) CTCF and cohesin regulate chromatin loop stability with distinct dynamics. eLife 6:e25776.

Heidemann M, Eick D. (2012) Tyrosine-1 and threonine-4 phosphorylation marks complete the RNA polymerase II CTD phospho-code. RNA Biol. 9:1144–6.

Heinz S, Texari L, Hayes MGB, et al. (2018) Transcription elongation can affect genome 3D structure. Cell 174:1522–1536.

Hsieh TS, Cattoglio C, Slobodyanyuk E, et al. (2020) Resolving the 3D landscape of transcription-linked mammalian chromatin folding. Mol Cell 78:539–553.

Hsiung CC, Bartman CR, Huang P, et al. (2016) A hyperactive transcriptional state marks genome reactivation at the mitosis-G1 transition. Genes Dev. 30:1423–1439.

Hug CB, Grimaldi AG, Kruse K, Vaquerizas JM. (2017) Chromatin architecture emerges during zygotic genome activation independent of transcription. Cell 169:216–228.

Ibrahim DM, Mundlos S. (2020) The role of 3D chromatin domains in gene regulation: a multi-facetted view on genome organization. Curr Opin Genet Dev. 61:1–8.

Jiang Y, Huang J, Lun K, et al. (2020) Genome-wide analyses of chromatin interactions after the loss of Pol I, Pol II, and Pol III. Genome Biol. 21:158.

Kim S, Shendure J. (2019) Mechanisms of interplay between transcription factors and the 3D genome. Mol Cell 76:306–319.

Kim Y, Shi Z, Zhang H, Finkelstein IJ, Yu H. (2019) Human cohesin compacts DNA by loop extrusion. Science 366:1345–1349.

Krietenstein N, Abraham S, Venev SV, et al. (2020) Ultrastructural details of mammalian chromosome architecture. Mol Cell 78:554–565.

Li Y, Haarhuis JHI, Sedeño Cacciatore Á, et al. (2020) The structural basis for cohesin-CTCF-anchored loops. Nature 578:472–476.

Marchal C, Sima J, Gilbert DM. (2019) Control of DNA replication timing in the 3D genome. Nat Rev Mol Cell Biol. 20:721–737.

Nagashima R, Hibino K, Ashwin SS, et al. (2019) Single nucleosome imaging reveals loose genome chromatin networks via active RNA polymerase II. J Cell Biol. 218:1511–1530.

Nielsen CF, Zhang T, Barisic M, Kalitsis P, Hudson DF. (2020) Topoisomerase IIα is essential for maintenance of mitotic chromosome structure. Proc Natl Acad Sci USA 117:12131–12142.

Nitiss JL. (2009) DNA topoisomerase II and its growing repertoire of biological functions. Nat Rev Cancer 9:327–37.

Nora EP, Goloborodko A, Valton AL, et al. (2017) Targeted degradation of CTCF decouples local insulation of chromosome domains from genomic compartmentalization. Cell 169:930–944.

Nuebler J, Fudenberg G, Imakaev M, Abdennur N, Mirny LA. (2018) Chromatin organization by an interplay of loop extrusion and compartmental segregation. Proc Natl Acad Sci USA 115:E6697–E6706.

Papantonis A, Cook PR. (2011) Fixing the model for transcription: the DNA moves, not the polymerase. Transcription 2:41–44.

Rada-Iglesias A, Grosveld FG, Papantonis A. (2018) Forces driving the three-dimensional folding of eukaryotic genomes. Mol Syst Biol. 14:e8214.

Rao SS, Huntley MH, Durand NC, et al. (2014) A 3D map of the human genome at kilobase resolution reveals principles of chromatin looping. Cell 159:1665–1680..

Rao SSP, Huang SC, Glenn St Hilaire B, et al. (2017) Cohesin loss eliminates all loop domains. Cell 171:305–320.

Rhodes JDP, Feldmann A, Hernández-Rodríguez B, et al. (2020) Cohesin disrupts polycomb-dependent chromosome interactions in embryonic stem cells. Cell Rep. 30:820–835.

Rosencrance CD, Ammouri HN, Yu Q, et al. Chromatin hyperacetylation impacts chromosome folding by forming a nuclear subcompartment. Mol Cell 2020; 78(1):112–126.

Rowley MJ, Lyu X, Rana V, et al. (2019) Condensin II counteracts cohesin and RNA polymerase II in the establishment of 3D chromatin organization. Cell Rep. 26:2890–2903.

Rowley MJ, Nichols MH, Lyu X, et al. (2017) Evolutionarily conserved principles predict 3D chromatin organization. Mol Cell 67:837–852.

Sati S, Bonev B, Szabo Q, et al. (2020) 4D genome rewiring during oncogene-induced and replicative senescence. Mol Cell 78:522–538.

Schwarzer W, Abdennur N, Goloborodko A, et al. (2017) Two independent modes of chromatin organization revealed by cohesin removal. Nature 551:51–56.

Singh S, Szlachta K, Manukyan A, et al. (2020) Pausing sites of RNA polymerase II on actively transcribed genes are enriched in DNA double-stranded breaks. J Biol Chem. 295:3990–4000.

Teves SS, An L, Bhargava-Shah A, Xie L, Darzacq X, Tjian R. (2018) A stable mode of bookmarking by TBP recruits RNA polymerase II to mitotic chromosomes. eLlife 7:e35621.

Thakurela S, Garding A, Jung J, Schübeler D, Burger L, Tiwari VK. (2013) Gene regulation and priming by topoisomerase IIα in embryonic stem cells. Nat Commun. 4:2478.

Thiecke MJ, Wutz G, Muhar M, et al. (2020) Cohesin-dependent and - independent mechanisms mediate chromosomal contacts between promoters and enhancers. Cell Rep. 32:107929.

Übelmesser N, Papantonis A. (2019) Technologies to study spatial genome organization: beyond 3C. Brief Funct Genomics 18:395–401.

Ulianov SV, Khrameeva EE, Gavrilov AA, et al. (2016) Active chromatin and transcription play a key role in chromosome partitioning into topologically associating domains. Genome Res. 26:70–84.

Uusküla-Reimand L, Hou H, Samavarchi-Tehrani P, et al. (2016) Topoisomerase II beta interacts with cohesin and CTCF at topological domain borders. Genome Biol. 17:182.

Vian L, Pękowska A, Rao SSP, et al. (2018) The energetics and physiological impact of cohesin extrusion. Cell 173:1165–1178.

Wang Y, Lu JJ, He L, Yu Q. (2011) Triptolide (TPL) inhibits global transcription by inducing proteasome-dependent degradation of RNA polymerase II (Pol II). PLoS One 6:e23993.

Wutz G, Ladurner R, St Hilaire BG, et al. (2020) ESCO1 and CTCF enable formation of long chromatin loops by protecting cohesinSTAG1 from WAPL. eLife 9:e52091.

Yesbolatova A, Natsume T, Hayashi KI, Kanemaki MT. (2019) Generation of conditional auxin-inducible degron (AID) cells and tight control of degron-fused proteins using the degradation inhibitor auxinole. Methods 164-165:73–80.

Zhang H, Emerson DJ, Gilgenast TG, et al. (2019) Chromatin structure dynamics during the mitosis-to-G1 phase transition. Nature 576:158–162.

Zirkel A, Nikolic M, Sofiadis K, et al. (2018) HMGB2 loss upon senescence entry disrupts genomic organization and induces CTCF clustering across cell types. Mol Cell 70:730–744.

Zuin J, Franke V, van Ijcken WF, et al. (2014) A cohesin-independent role for NIPBL at promoters provides insights in CdLS. PLoS Genet. 10:e1004153.

## REFERENCES FOR METHODS

Abraham BJ, Hnisz D, Weintraub AS, et al. (2017) Small genomic insertions form enhancers that misregulate oncogenes. Nat Commun. 8:14385.

Arrigoni L, Richter AS, Betancourt E, et al. (2016) Standardizing chromatin research: a simple and universal method for ChIP-seq. Nucleic Acids Res. 44:e67.

Buenrostro JD, Wu B, Chang HY, Greenleaf WJ. (2015) ATAC-Seq: A method for assaying chromatin accessibility genome-wide. Curr Protoc Mol Biol. 109: 21.29.1–21.29.9.

Crane E, Bian Q, McCord R, et al. (2015) Condensin-driven remodelling of X chromosome topology during dosage compensation. Nature 523:240–244.

Corces RM, Trevino AE, Hamilton EG, et al. (2017) An improved ATAC-seq protocol reduces background and enables interrogation of frozen tissues. Nat Methods 14:959–962.

Dobin A, Davis CA, Schlesinger F, et al. (2013) STAR: ultrafast universal RNA-seq aligner. Bioinformatics 29:15–21.

Durand NC, Shamim MS, Machol I, et al. (2016) Juicer Provides a One-Click System for Analyzing Loop-Resolution Hi-C Experiments. Cell Syst. 3:95–98.

Greenwald WW, Li H, Smith EN, et al. (2017) Pgltools: a genomic arithmetic tool suite for manipulation of Hi-C peak and other chromatin interaction data. BMC Bioinformatics 18:207.

Kharchenko PV, Tolstorukov MY, Park PJ. (2008) Design and analysis of ChIP-Seq experiments for DNA-binding proteins. Nat Biotech. 26:1351–1359.

Langmead B and Salzberg SL. (2012) Fast gapped-read alignment with Bowtie 2. Nat Methods 9:357–359.

Li H, Handsaker B, Wysoker A, et al. (2009) The sequence alignment/map format and SAMtools. Bioinformatics 25:2078–2079.

Li Q, Brown JB, Huang H, Bickel PJ. (2011) Measuring reproducibility of high-throughput experiments. Ann. Appl. Stat. 5:1752–1779.

Liao Y, Smyth GK, Shi W. (2014) FeatureCounts: an efficient general purpose program for assigning sequence reads to genomic features. Bioinformatics 30:923–930.

Love MI, Huber W, Anders S. (2014) Moderated estimation of fold change and dispersion for RNA-seq data with DESeq2. Genome Bio. 15:550.

Marx A, Backes C, Meese E, Lenhof HP, Keller A. (2016) EDISON-WMW: Exact dynamic programing solution of the Wilcoxon-Mann-Whitney test. Genom Proteom Bioinform. 14:55–61.

Melnik S, Caudron-Herger M, Brant L, Carr IM, Rippe K, Cook PR, Papantonis A. (2016) Isolation of the protein and RNA content of active sites of transcription from mammalian cells. Nat Protoc. 11:553–565.

Plimpton S. Fast parallel algorithms for short-range molecular dynamics. (1995) J Comp Phys. 1:1–19.

Ramírez F, Dündar F, Diehl S, Grüning BA, Manke T. (2014) DeepTools: A flexible platform for exploring deep-sequencing data. Nucleic Acids Res. 42:W187–91.

Risso D, Ngai J, Speed TP, Dudoit S. (2014) Normalization of RNA-seq data using factor analysis of control genes or samples. Nat Biotech. 32:896–902.

Roukos V, Pegoraro G, Voss TC, Misteli T. (2015) Cell cycle staging of individual cells by fluorescence microscopy. Nat Protoc. 10:334–348.

Rowley MJ, Poulet A, Nichols MH, et al. (2020) Analysis of Hi-C data using SIP effectively identifies loops in organisms from *C. elegans* to mammals. Genome Res. 30:447–458.

Subramanian A, Tamayo P, Mootha VK, et al. (2005) Gene Set Enrichment Analysis: A Knowledge-Based Approach for Interpreting Genome-Wide Expression Profiles. Proc Natl Acad Sci USA 102:15545.

Watrin E, Schleiffer A, Tanaka K, et al. (2006) Human Scc4 is required for cohesin binding to chromatin, sister-chromatid cohesion, and mitotic progression. Curr Biol. 16:863–874.

Zhang Y, Liu T, Meyer CA, et al. (2008) Model-based analysis of ChIP-seq (MACS). Genome Biol. 9:R137.

